# A framework for the quantification of soundscape diversity using Hill numbers

**DOI:** 10.1101/2022.01.11.475919

**Authors:** Thomas Luypaert, Anderson S. Bueno, Gabriel S. Masseli, Igor L. Kaefer, Marconi Campos-Cerqueira, Carlos A. Peres, Torbjørn Haugaasen

## Abstract

1. Soundscape studies are increasingly common to capture landscape-scale ecological patterns. Yet, several aspects of soundscape diversity quantification remain unexplored. Although some processes influencing acoustic niche usage may operate in the 24h domain, most acoustic indices only capture the diversity of sounds co-occurring in sound files at a specific time of day. Moreover, many indices do not consider the relationship between the spectral and temporal traits of sounds simultaneously. To provide novel insights into landscape-scale patterns of acoustic niche usage at broader temporal scales, we present a workflow to quantify soundscape diversity through the lens of functional ecology.
2. Our workflow quantifies the functional diversity of sound in the 24-hour acoustic trait space. We put forward an entity, the Operational Sound Unit (OSU), which groups sounds by their shared functional properties. Using OSUs as our unit of diversity measurement, and building on the framework of Hill numbers, we propose three metrics that capture different aspects of acoustic trait space usage: (i) soundscape richness; (ii) soundscape diversity; (iii) soundscape evenness. We demonstrate the use of these metrics by (a) simulating soundscapes to assess if the indices possess a set of desirable behaviours; and (b) quantifying the soundscape richness and evenness along a gradient in species richness to illustrate how these metrics can be used to shed unique insights into patterns of acoustic niche usage.
3. We demonstrate that: (a) the indices outlined herein have desirable behaviours; and (b) the soundscape richness and evenness are positively correlated with the richness of soniferous species. This suggests that the acoustic niche space is more filled where taxonomic richness is higher. Moreover, species-poor acoustic communities have a higher proportion of rare sounds and use the acoustic space less effectively. As the correlation between the soundscape and taxonomic richness is strong (>0.8) and holds at low sampling intensities, soundscape richness could serve as a proxy for taxonomic richness.
4. Quantifying the soundscape diversity through the lens of functional ecology using the analytical framework of Hill numbers generates novel insights into acoustic niche usage at a landscape scale and provides a useful proxy for taxonomic richness measurement.

## 1. Introduction

The recent emergence of Passive Acoustic Monitoring (PAM) offers promising opportunities for ecological monitoring. Automated acoustic sensors can make recordings of environmental sound at broad spatio-temporal scales with reduced cost and human effort compared to equivalent active acoustic sampling by an *in-situ* observer (Gibb et al. 2019). Using these acoustic data, the taxonomic diversity of a biological community can be derived by isolating and identifying species’ calls, thus providing an objective and permanent record of the resident soniferous biological community (e.g., birds, mammals, amphibians, and insects; Gibb et al. 2019; Sugai et al. 2019). Yet, obtaining species-level information for broad spatio-temporal scales or wide taxonomic breadth presents numerous logistical and analytical difficulties, such as the time-consuming and knowledge-demanding nature of aural annotation, and the paucity of reliable automated species identifiers and reference databases for most taxa and regions (Toledo et al. 2015; Gibb et al. 2019; Sugai et al. 2019).

In addition to taxonomic information, species’ sounds carry functional significance. They are essential for a range of social interactions, including courting behaviour, territorial defence, predator avoidance, and food sharing (Darwin 1872; Seyfarth and Cheney 2003). As such, species’ sounds are subject to selective pressures at multiple scales (Zsebők et al. 2021), resulting in a staggering variety of acoustic functional traits that are expressed in the timing, frequency and amplitude features of the acoustic signal. The field of soundscape ecology exploits this variation in acoustic traits, attempting to infer ecological information from the soundscape – i.e. the collection of biological (biophony), geophysical (geophony) and human-produced (anthrophony) sounds emanating from a landscape – without the need for species identification (Krause 1987; Pijanowski et al. 2011a; Pijanowski et al. 2011b). This discipline builds on the premise that the diversity of acoustic traits in the landscape can be used to understand ecological processes and nature-human dynamics across spatial and temporal scales (Pijanowski et al. 2011b). To accomplish this, more than 60 acoustic indices have been developed to date (Buxton et al. 2018), each of which captures the diversity of acoustic functional traits across the time-frequency domain of a sound file in some way.

The diversity of acoustic signals in functional trait space can be used to shed light on underlying ecological and evolutionary mechanisms, such as processes influencing acoustic species assembly (Gasc et al. 2013). For instance, one of the cornerstone theories of soundscape ecology is the Acoustic Niche Hypothesis, which states that acoustic space is a core ecological resource for which soniferous sympatric species compete, leading to the partitioning of the acoustic niche in the time-frequency domain to avoid spectro-temporal overlap in sound production (Krause 1993). Following this reasoning, a more speciose community should lead to increased competition and partitioning of acoustic niche space, which can be measured by quantifying the functional diversity of sounds in the acoustic trait space. Indeed, acoustic indices have been successfully applied as proxies for the diversity of species (Depraetere et al. 2012; Towsey et al. 2014) or sound types (Pijanowski et al. 2011b) in the corresponding sound file. Moreover, they have also been used as descriptors of landscape configuration and ecosystem health (Tucker et al. 2014; Fuller et al. 2015), diel patterns in different environments (Rodriguez et al. 2014; Farina et al. 2015), seasonal changes in soundscapes (Pieretti et al. 2011), habitat identity (Pijanowski et al. 2011b; Depraetere et al. 2012), among others. However, despite the link to niche theory and acoustic functional trait measurement, the field of soundscape ecology has never been explicitly considered through the lens of functional ecology, thereby potentially precluding insights into acoustic community assembly processes at landscape scales.

We argue that several aspects of soundscape diversity quantification remain unexplored. For instance, most indices capture acoustic patterns using either time-averaged spectrograms (collapsed in the temporal domain) or measures of variation in amplitude over time (collapsed in the frequency domain). As highlighted by Eldridge et al. (2016), these indices are fundamentally limited in their ability to detect functional diversity patterns across both the spectral and temporal dimensions simultaneously. Since spectro-temporal partitioning of the acoustic niche might be one of the processes dictating acoustic community assembly, considering both the spectral and temporal dimensions of the acoustic trait space simultaneously may be key to evaluating how the acoustic niche is structured. Moreover, most existing acoustic indices are calculated over relatively short-duration timescales (e.g., 1 minute sound files). Here, we posit that the assembly processes that structure the presence and distribution of sound in the acoustic functional trait space should also be considered at broader temporal scales. As many species repeat their sound emissions at circadian timescales (Agostino et al. 2020), some of the temporal partitioning of the acoustic niche likely occurs in the 24-hour time domain. As such, the diversity of all acoustic signals occurring in these 24 hours should be considered in relation to each other. Yet, for the field of soundscape ecology, explicit quantification of the functional relationship between sounds in the 24-hour acoustic trait space at a landscape scale is scarce (but see Aide et al. 2017). To do so, we require a robust framework that produces informative metrics that can capture within- and between-soundscape differences in spectro-temporal trait space usage.

In this work, we describe a workflow to decompose the functional diversity of sound in the acoustic trait space, hereafter referred to as the soundscape diversity. This workflow is grounded in the principles of acoustic niche theory and leans heavily on the field of functional ecology. Rather than focussing on the fine-scale temporal patterns (*i.e.*, bioacoustics studies) or assessing the soundscape diversity of an acoustic assemblage at a particular time of day (*i.e.*, many soundscape studies), we propose a framework to investigate the functional relationship between all sounds produced in the broader 24h spectro-temporal functional trait space at a given geographic location. To do so, we put forward a novel entity, the Operational Sound Unit (OSU), which groups sounds by their shared functional properties in the acoustic trait space. Using OSUs as our unit of soundscape diversity measurement, and building on the framework of Hill numbers, we propose three metrics that capture different aspects of the functional diversity of the soundscape: (i) soundscape richness; (ii) soundscape diversity; (iii) soundscape evenness.

Our workflow offers some unique insights to complement existing soundscape diversity metrics. For instance, dissecting the soundscape diversity into its facets can provide insights into various aspects of 24-hour acoustic trait space usage, including patterns of niche saturation, evenness, dominance, or rarity. Moreover, using the analytical framework of Hill numbers, we can quantify the soundscape diversity at various scales, decomposing the regional metacommunity diversity (γ-diversity) into its local diversities (α-diversity) and a community turnover component (β-diversity) using a simple multiplicative relationship. Additionally, Hill numbers can also be used to quantify taxonomic, functional, and phylogenetic diversity, which ensures that observed relationships between soundscape diversity and other facets of biodiversity represent real-world ecological patterns rather than mathematical artefacts that stem from different formulae. If the Acoustic Niche Hypothesis holds, this means the various soundscape diversity components could be used to shed light on the taxonomic richness or diversity of the local soniferous community using a common framework of reference.

To illustrate these unique benefits, we show that the proposed soundscape diversity metrics follow a set of fundamental criteria for functional diversity metrics and act in an ecologically intuitive way. Moreover, using an acoustic dataset from Brazilian Amazonia, we investigate how the soundscape diversity metrics behave along an ecological gradient in species richness. We report positive correlations for both the soundscape richness and soundscape evenness with the richness of soniferous species in an acoustically complex rainforest setting.

## 2. Methods

The implementation of this workflow is facilitated by the ‘soundscapeR’ package, written in the R-programming language (R Core Team 2020) and currently under development on GitHub (https://github.com/ThomasLuypaert/soundscapeR).

### 2.1. Defining the acoustic trait space

Before quantifying the functional diversity of a system, the functional trait space in which units will be compared needs to be defined. For this, the functional variables under consideration largely determine the functional trait space they define (Zsebők et al. 2021). Regardless of which processes influence acoustic community assembly, the timing, frequency, and amplitude features of sounds are important functional traits of acoustic signals that are subject to evolutionary processes. As such, we use the timing and frequency of sounds as the variables that delineate a two-dimensional acoustic trait space and employ an amplitude-based threshold value to quantify the detection/non-detection of sounds within this acoustic space.

Although soniferous species are known to produce sounds from infrasound to deep into the ultrasonic spectrum, mostly for bats and katydids, we recommend constraining the upper frequency limit to 22,050 Hz, approximately the maximal frequency audible to humans (Farina 2013). We employ this strategy for several reasons. Most wildlife sounds can be found in this frequency range (Farina and James 2016), so the evolutionary mechanisms thought to structure acoustic assemblages are likely strongest in this range. Moreover, in downstream analyses, we employ spectral acoustic indices to capture the amplitude structure of sounds. As ultrasonic frequencies are usually omitted from soundscape studies employing acoustic indices, we cannot guarantee the sensitivity of these indices to the amplitude features of ultrasonic sounds. We therefore omit these frequencies from the trait space definition.

In the temporal domain, we follow Aide et al. (2017) and consider the temporal dimension of acoustic trait space as the 24h period in which species emit sounds, rather than focussing on the fine-scale temporal structure of signals that is often the subject of inquiry in bioacoustics studies (Zsebők et al. 2021). The reasoning here is twofold. First, we are interested in investigating the relationship between all sounds produced at a given site, not just between the sounds of a site’s acoustic assemblage producing sound at a particular time. Second, almost all living organisms have a circadian timing system that allows physiological and behavioural adjustments to the 24h cyclic variation in the surrounding environment (Agostino et al. 2020). Indeed, many soniferous species have been shown to repeat their sound emissions at circadian timescales (Jianguo et al. 2011; Wang et al. 2012; Da Silva et al. 2014). Therefore, a 24h period seems like an appropriate sample duration to capture and compare most resident species’ sounds at any given site.

### 2.2. Defining an entity for soundscape diversity measurement

The next step in quantifying the functional diversity of a system is to define an entity of measurement. In functional ecology, diversity metrics are usually based on the functional traits of taxonomic species and their abundance. Yet, taxonomic information is not always available. In some fields of research where the taxonomic identity of individuals is unknown, Operational Taxonomic Units (OTUs) - or groups of related individuals which share a set of observed characteristics (Sokal and Sneath 1963) - are widely accepted as a base entity to infer the diversity of a system. Here, we attempt to measure and compare the functional properties of entities (sounds) in a system (functional trait space) without a taxonomic link to the source organisms. Hence, to quantify the functional diversity of sounds in acoustic trait space (hereafter referred to as soundscape diversity), we require an entity that groups sounds by their shared functional properties without the need for taxonomic information.

According to Mouchet et al. (2010), measuring the functional diversity of a system is equivalent to quantifying the presence and distribution of functional units in multidimensional space, and functionally identical units should be grouped into one. In analogy to OTUs for taxonomic diversity studies, we therefore propose a new type of entity - the Operational Sound Unit (OSU) - which groups sounds by their shared spectro-temporal properties. We use this entity as the base unit by which to measure the various properties of soundscape diversity. OSUs are obtained by subdividing the acoustic trait space into many discrete spectro-temporal bins that can be seen as the soundscape equivalent of the time-frequency bins in a spectrogram. Despite being conceptually analogous to the time-frequency bins used to calculate the ‘Acoustic Space Use’ (ASU) metric described in Aide et al. (2017), the OSU differs in the amplitude features that are used to capture the presence and abundance of sound in acoustic trait space, and in the resolution along the temporal axis, which will be further clarified in the next section.

#### 2.2.1. Assessing the presence of sound in acoustic trait space

Before assessing the soundscape diversity, acoustic data needs to be collected. For this, several decisions need to be made. First, the sampling rate and bit depth of the acoustic recorder need to be chosen. These parameters will dictate the frequency and amplitude resolution. The sampling rate should be twice the desired maximal frequency, a principle known as the Nyquist-Shannon sampling theorem. In this study, the sampling rate needs to be a minimum of 44,100, as we are interested in sounds up to 22,050 Hz. The choice of sampling rate and bit depth also constitutes a trade-off between the desired resolution on the one hand, and the available storage on the memory card and the battery life of the sensor on the other hand, as higher sampling rates and bit depths are more storage- and energy-demanding. Next, the recording schedule and duration for each site should be determined. For soundscape studies, sound files are usually collected for 1-min durations. If multiple soundscapes are to be compared in the same study, the same recording schedule should be used. The soundscape can be recorded either continuously or using a regular-interval sampling regime (1 min/5 min; 10 min/30 min; etc.). However, sparse sampling regimes are generally discouraged as they require the soundscape to be recorded for long periods before the soundscape variability is captured adequately, which in turn introduces issues related to seasonal variation (Bradfer-Lawrence et al. 2019). In S.1, we provide recommendations regarding the choice of sampling duration and regime using our workflow.

Once the acoustic data is acquired, we start by quantifying the presence of sound with specific spectro-temporal amplitude features in the acoustic trait space during the recording period. The amplitude features of interest will determine how the acoustic trait space is defined and will characterise different aspects of soundscape diversity. Here, we are interested in all biological sounds produced at a given site, regardless of which source they emanate from. We therefore focus on the presence of sounds exceeding an amplitude threshold for a certain duration of time in each 1-min recording.

As introduced in section 2.1, we consider the temporal dimension of acoustic trait space as the 24h period in which species produce sound. We pool sound files in the recording period of a specific site into 24h samples of the acoustic trait space, each sample containing the sound files obtained in a single day (00:00h – 23:59h; Fig. 1A). To determine where (frequency domain) and when (time-domain) in the acoustic space sound is present throughout the recording period, we rely on the properties of the Acoustic Cover (CVR) spectral acoustic index. For each 1-minute sound file, the CVR index produces a vector of N-values, one for each frequency bin of the spectrogram, and calculates the proportion of cells in each noise-reduced frequency bin that exceed a 3 dB amplitude value, resulting in a value ranging between 0 and 1 (consult Towsey (2017) for a detailed breakdown of index computation). We calculate the CVR index for all 1-min sound files in each 24h sample. Acoustic recordings are processed following the methods outlined in Towsey (2017), computing indices using QUT Ecoacoustics Analysis Programs software (Towsey et al. 2018; Fig. 1B).

**Figure 1:**
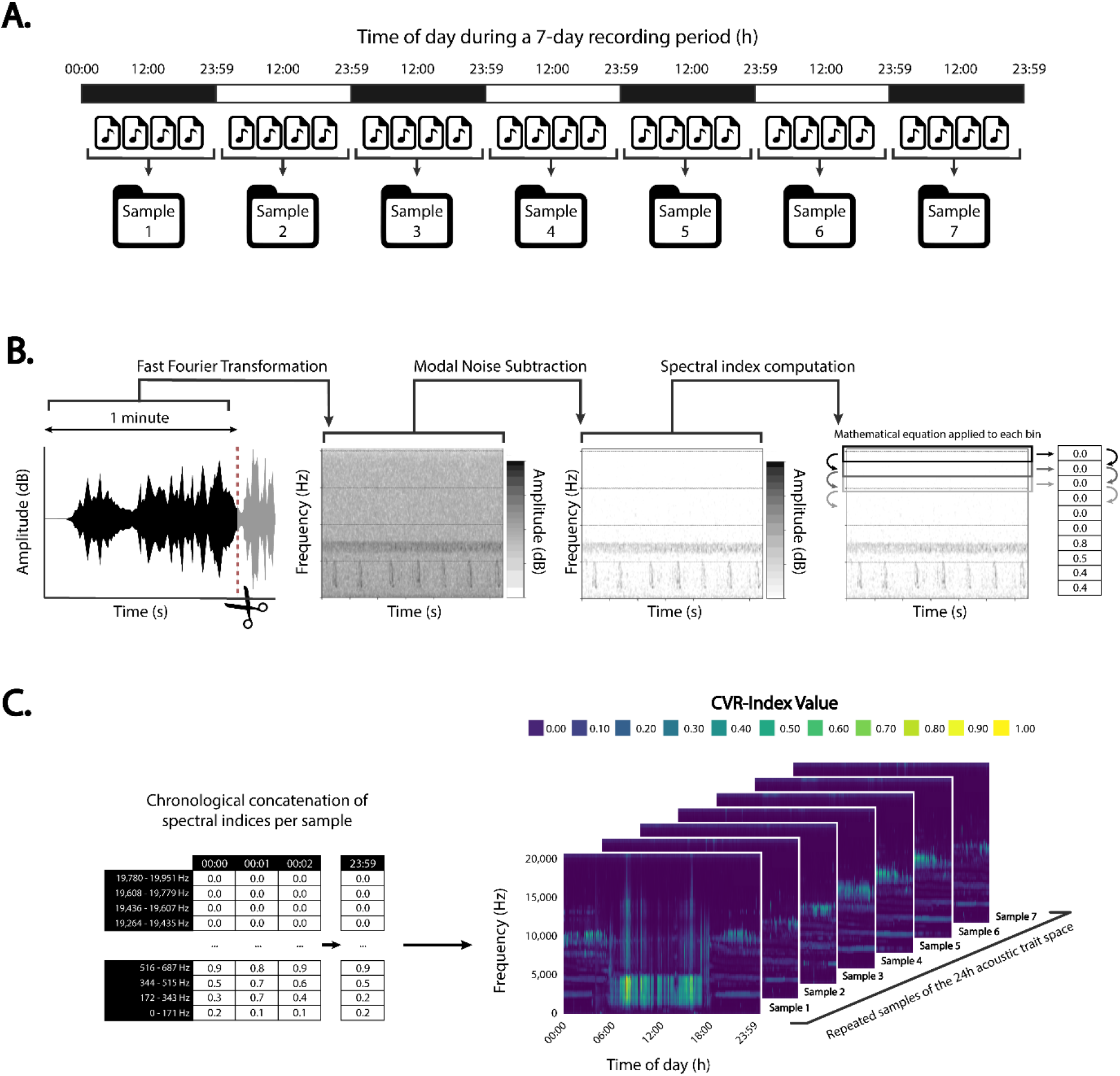
A visual representation of the workflow steps to assess the presence of sound in acoustic trait space. **A.** Sounds in the recording period (7 days) are pooled into 24h samples of the acoustic trait space. **B.** For each sample, all sound files are cut into 1-min segments. Each 1-min segment is subjected to a Fast Fourier Transformation, followed by Model Noise Subtraction and Spectral Index Computation - resulting in a spectral index vector (CVR-index) for each 1-min file. **C.** For all sound files per sample, the CVR-index vectors are concatenated chronologically, resulting in a data frame with time-of-day as columns, frequency bins as rows, and the CVR-value as cells. Finally, we obtain repeated samples of the 24h acoustic trait space, each of which showing the presence of sound in the time-frequency domain.

Once the CVR-index has been calculated for the sound files in each 24h sample, the spectral index vectors for all 1-min files in a sample are concatenated chronologically, creating a data frame with the time of recording as columns, the frequency bins as rows, and the value of the CVR index for each time-frequency bin as cells. This provides a view of the presence and distribution of sound in each sample of the 24h acoustic trait space (Fig. 1C).

#### 2.2.2. Operational Sound Units (OSUs)

By assessing the presence of sound in acoustic trait space as described in section 2.2.1, we have divided the trait space into discrete time-frequency bins that group sounds by their shared functional properties (shared time and frequency values in trait space), thus capturing our concept of Operational Sound Units (Fig. 2).

**Figure 2:**
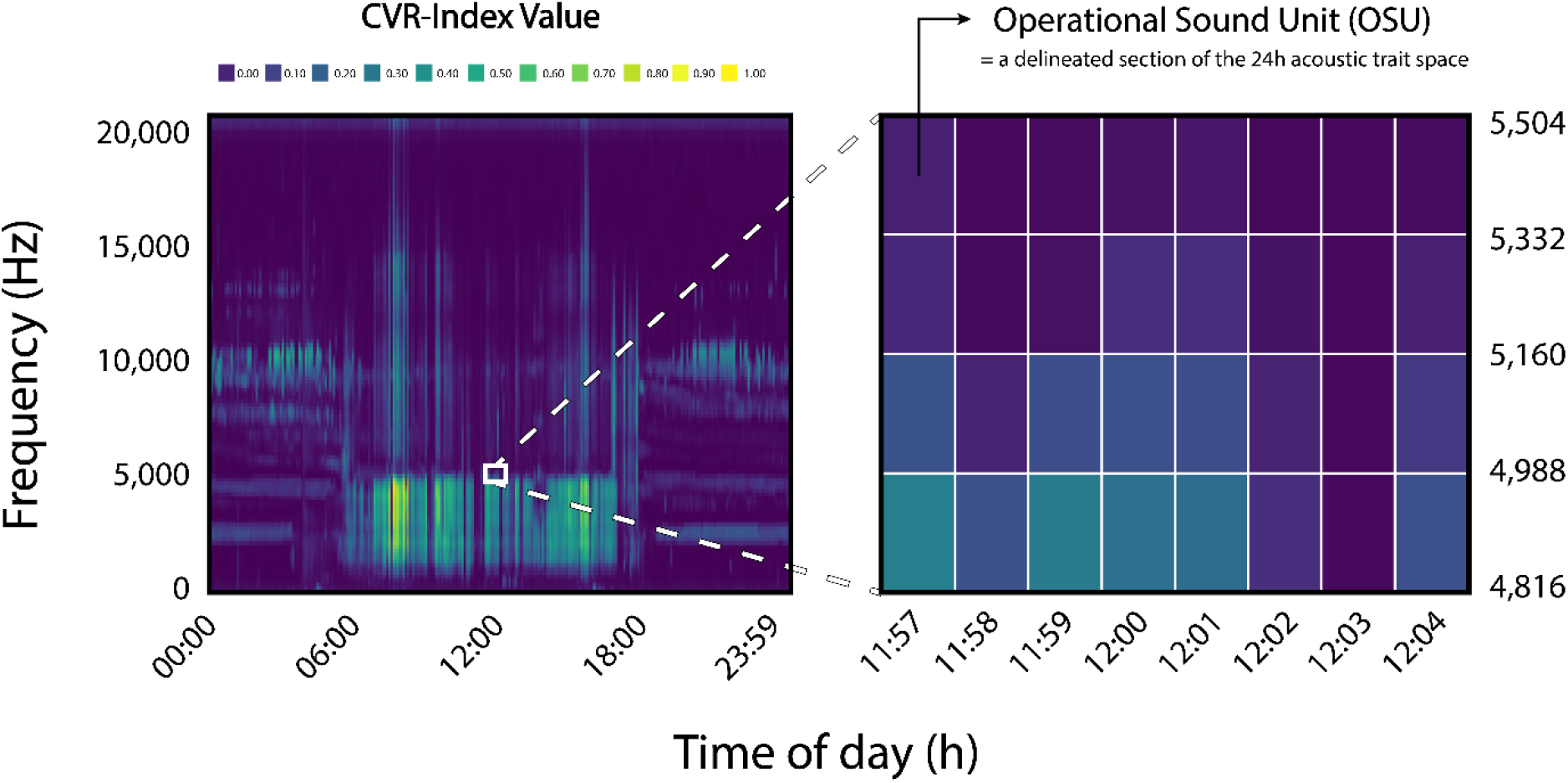
A conceptual visualisation of Operational Sound Units (OSUs) in the 24h acoustic trait space. Each 24h sample of the acoustic trait space can be divided into sections which we define as Operational Sound Units (OSUs). These OSUs are delineated by the frequency-bin width of the spectral index vector (frequency domain) and the recording interval of the sampling regime (temporal domain), and group sounds by their shared functional properties in acoustic space.

As with time-frequency bins in spectrograms, the resolution of OSUs in acoustic trait space, and thus the total number of OSU in which the space is partitioned, is variable. In the temporal domain, the width of OSUs and the total number of OSUs in trait space is dictated by the file length for index calculation and the sampling regime of the study (continuous vs regular interval), respectively. The 1-minute duration employed for index calculation retains enough detail in the acoustic features for long-duration soundscape analysis, facilitates rapid computation, and has been used as the *de facto* standard in most soundscape studies (Truskinger and Towsey 2019). For the sampling regime, sparse regimes are generally discouraged (but see S.1 for further details). In this study, we use a 1/5 min sampling regime (see section 3).

In the frequency domain, the resolution of OSUs is determined by the width of the frequency bins of the spectral index vectors. This is in turn dictated by the sampling rate and window length, which are specified in the short-time Fourier transformation. As mentioned earlier, we truncate the frequency domain for this workflow at 22,050 Hz and thus require a sampling rate of 44,100. The choice of window length will therefore determine the OSU resolution of the frequency domain. Choosing the appropriate window length will depend on the soniferous community of interest. In S.2, we provide guidance on window length choice, and recommend using a 256-sample window length.

Considering the aforementioned settings (44,100 Hz sampling rate, 1/5 min sampling regime and a window length of 256), the frequency domain consists of 128 frequency bins (number of bins = window length / 2) of 172 Hz width (bin width = (sampling rate / 2) / number of bins). The temporal domain consists of 288 bins (24 hours = 1440 min with 1/5 min recorded = 288 bins). As such, the total number of detectable OSUs in the trait space using these settings is 36,864 (128 frequency bins * 288 temporal bins).

### 2.3. Assessing the prevalence of OSUs in acoustic trait space

As mentioned in section 2.2., functional diversity is typically quantified using the diversity of species traits and a measure of species importance such as abundance (Chiu and Chao 2014). Up to this point, we have defined an entity of measurement (OSU) and quantified its amplitude features in acoustic trait space across the recording period. Next, we need to attribute an importance value to each OSU.

Instead of using the raw CVR-values obtained in section 2.2.1, we opt to use an incidence-based approach to derive an importance value for each OSU. We quantify the detection/non-detection of OSUs with specific amplitude features in each 24h sample of acoustic trait space. As the CVR values used in this study rarely take zero values, we convert the CVR index values to a binary variable representing detection (1) or non-detection (0) using a threshold (Fig. 3A). The choice of the threshold will depend on the study system and is influenced by sound transmission characteristics of the habitat and the amount of ambient noise in the surrounding environment (Darras et al. 2016). For a comparison of thresholding methods, consult S.3. Regardless of which binarisation method is adopted, we recommend visually inspecting the pre- and post-binarisation soundscapes to ensure the amplitude structure is captured without removing excess sound. Once the detection/non-detection of OSUs in each 24h sample of acoustic trait space has been established, we compute the relative OSU abundance in acoustic trait space. For this, we add up the number of times each OSU was detected (value = 1) across all 24h samples in the recording period and divide this by the number of 24h samples to obtain the incidence frequency (Figs. 3B-C). To avoid confusion between the sound frequency (Hz) and the incidence frequency (relative number of OSU occurrences), we will henceforth refer to the OSU importance value as the relative abundance.

**Figure 3:**
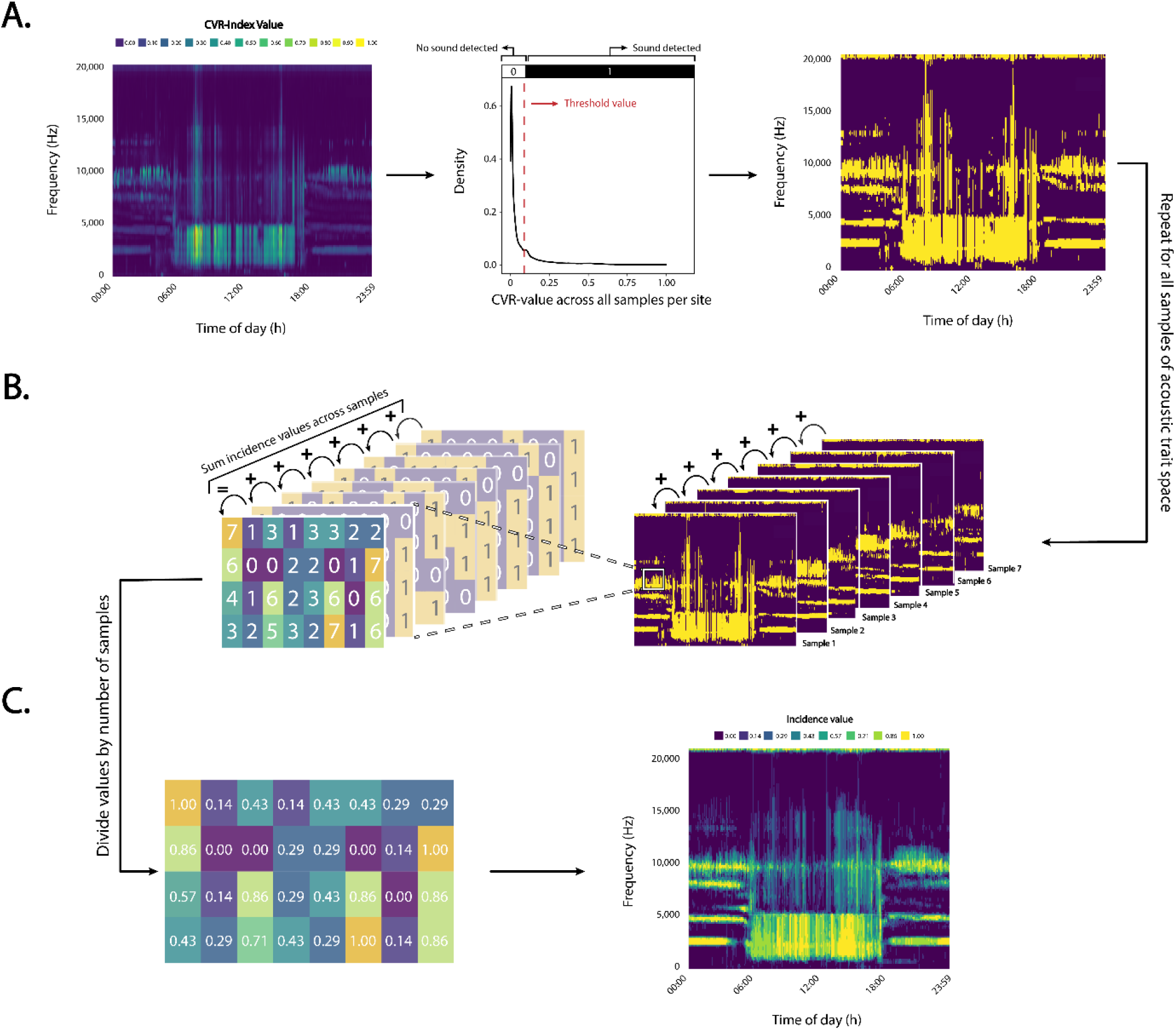
A conceptual representation of the methodology used to attribute an importance value to OSUs in acoustic trait space. **A.** A binarisation algorithm is applied to each sample of acoustic trait space, resulting in a binary variable representing the detection / non-detection of OSUs across samples; **B.** For each OSU, the detection (1) / non-detection (0) values are summed across all 24h samples of acoustic trait space and divided by the number of samples to obtain the OSU relative abundance (incidence frequency); **C.** The presence, relative abundance, and distribution of OSUs in acoustic trait space.

### 2.4. Quantifying soundscape diversity using Hill numbers

Based on Mason et al. (2005) and Villéger et al. (2008), functional diversity can be broken down into three components: (i) functional richness; (ii) functional evenness; (iii) functional divergence / distinctiveness / dispersion. Here, we will focus on the quantification of soundscape richness and evenness, and add an additional component, the soundscape diversity, which incorporates components of the former two. Within functional ecology, a plethora of indices have been proposed to measure these various diversity components (see Pavoine and Bonsall (2011) for an overview). Rao’s quadratic entropy (Q) has been commonly applied to the quantification of functional diversity (Chao et al. 2019). Yet, like many entropy indices, Rao’s Q does not abide by the replication principle, which reduces its utility for comparative purposes and can yield ecologically unintuitive behaviours (Jost 2006; Chao et al. 2014).

Instead, Hill numbers appear to be the most appropriate framework for decomposing the diversity of a system into its various components (Hill 1973; Jost 2006; Chao et al. 2014). Unlike the entropy indices, Hill numbers scale proportionally with the underlying diversity – when the diversity of the system doubles, so does the value of the index (replication principle – see S.5.3 for demonstration). Moreover, the Hill numbers framework can be used to measure not only the soundscape diversity, but also the taxonomic, functional, and phylogenetic diversity, giving metrics a common behaviour, interpretation, and standardised unit. This ensures comparisons between the soundscape diversity and other diversity types represent real-world ecological patterns, rather than mathematical artefacts stemming from different formulae (Chao et al. 2014). Finally, this framework also allows decomposing the regional metacommunity diversity (γ-diversity) into its local diversity (α-diversity) and community turnover (β-diversity) components.

Hill numbers are computed as follows:

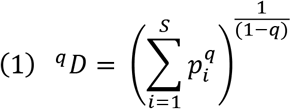

With S being the number of OSUs, *pi* the relative abundance of OSU *i*, and *q* the order of diversity. This equation expresses the diversity of the system as the ‘effective number of entities’ (OSUs) – the number of equally abundant OSUs that would yield the same value of diversity.

Here, we briefly describe the soundscape richness, diversity and evenness components and introduce the indices used for their measurement in the acoustic trait space.

#### Soundscape richness and diversity

The sensitivity to the relative abundance of OSUs is modulated using the order of diversity (*q*) without changing the interpretation of *^q^D*. When *q=0*, the relative abundance is disregarded and equation (1) yields *^0^D*=S, i.e., the richness of OSUs in acoustic trait space – or the soundscape richness. The functional richness generally quantifies the amount of acoustic trait space that is filled by the community (Mason et al. 2005). In our workflow, the soundscape richness measures the amount of acoustic trait space occupied by OSUs throughout the recording period without considering their relative abundance. Although being methodologically different, conceptually our functional soundscape richness metric is analogous to the soundscape saturation metric described in Burivalova et al. (2018), who measure the saturation of acoustic trait space at a 1-min scale. Similarly, our metric is related to the acoustic space use (ASU) metric described in Aide et al. (2017), which quantifies the saturation of acoustic trait space on a 24-hour scale, but uses a different methodology to detect sounds and aggregates those sounds at broader 1-hour intervals.

The higher the order of diversity *q*, the more weight is given to highly abundant OSUs. For instance, when *q*=1, the soundscape diversity *^1^D* equals the exponential of the Shannon entropy or the number of common OSUs in the soundscape. Similarly, when *q*=2, the soundscape diversity *^2^D* equals the reciprocal of the Simpson index, or the number of dominant or highly abundant OSUs in the soundscape. These three Hill numbers represent simple transformations of the traditional diversity indices which have already been used for decades and calculate mean species rarity using the arithmetic (q=0), geometric (q=1), and harmonic means (q=2) (Hill 1973). Although the soundscape richness and diversity metrics are usually expressed in the total number of OSUs, soundscape metrics can still be compared between soundscapes with differing dimensions (a different number of detectable OSUs due to window length / sampling regimes differences) by dividing the soundscape richness or diversity by the total number of detectable OSUs in the soundscape, thus generating the percentage of trait space filled with sound.

#### Soundscape evenness

The functional evenness describes the equitability of abundances in functional trait space (Mason et al. 2005). Following Jost (2010), various measures of evenness can be calculated by taking the ratio between Hill numbers *^q^D* with *q* = 1, 2,…, and the richness *^0^D*. Here, the choice of *q*-values determines the importance of OSU abundance on the evenness metric. For instance, since *^1^D* roughly represents the number of common OSUs in the acoustic trait space, the evenness ratio ^1^*D* / *^0^D* represents the proportion of common OSUs in the community. Similarly, as *^2^D* represents the number of dominant OSUs, the evenness ratio represents the proportion of dominant OSUs. Different *q*-values differ in the sharpness of the cut-off between rarity, commonness, or dominance. These patterns in evenness are best represented by constructing diversity profiles, a type of visualisation showing a series of Hill numbers derived using a continuous function of the order of diversity *q* (Jost 2007; Chao et al. 2012; see Fig. S11). Diversity profiles provide the most complete representation of the soundscape evenness, giving the relative abundance distribution of OSUs in the soundscape, and highlighting the change in diversity with changing importance of commonness or rarity. As the soundscape diversity and evenness can both be calculated for an infinite amount of q-values, for the remainder of this work we will define diversity as *^2^D* and evenness as *^2^D* / *^0^D* (Jost 2006). We make this choice because *q = 2* corresponds to a commonly used biodiversity metric used in literature (the reciprocal of the Simpson index), and the q-value is large enough to incorporate patterns of rarity and dominance in the acoustic community.

In S.4, we outline the theoretical framework for decomposing the soundscape diversity into its alpha, beta, and gamma components. Moreover, in S.5 we illustrate the behaviour and intuitive properties of the proposed soundscape diversity metrics by simulating artificial soundscapes. The simulated datasets serve to demonstrate the behaviour of the metrics with respect to some fundamental criteria for functional diversity metrics, as outlined in Ricotta (2005), Villéger et al. (2008) and Mouchet et al. (2010).

## 3. Case study

In this section, we explore the behaviour of our metrics of soundscape diversity in a real-life ecological setting. Specifically, we characterised the soundscape richness and evenness along a gradient in the richness of soniferous species using an empirical dataset from Brazilian Amazonia (1°40’S, 59°40’W). Under the Acoustic Niche Hypothesis, we expect the soundscape richness to be positively related to the richness of soniferous species (Krause 1993). For the soundscape evenness, we don not expect a relationship with taxonomic richness unless changing species richness was associated with a shift in the relative abundance distribution of the acoustic community (Wilsey et al. 2005).

### 3.1. Data collection

Acoustic data were collected at the Balbina Hydroelectric Reservoir (BHR) in Brazilian Amazonia. Surveys were conducted between July and December 2015, the local dry season, sampling 151 plots at 78 sampling sites (74 forest islands and 4 continuous forest sites) in the mainland (Bueno et al. 2020). At each plot, a single acoustic recorder was deployed at 1.5 m height and set to record the soundscape for 1-min every 5-min for 5-9 consecutive days at a sampling rate of 44.1 kHz using the ARBIMON Touch application. A subset of sites was taken for which previously published data on the taxonomic richness of anurans and primates was available (see section 3.2). For avian species, new taxonomic richness data for these forest islands was generated (see S.6.2). Moreover, several plots were omitted due to insufficient recording quality, persistent noise or positioning in riparian habitat. Ultimately, 65 plots located at 35 sampling sites (32 islands and three continuous forest sites) were retained (see S.6.1). If multiple plots were present at the same site, data were aggregated across plots, resulting in a final sample size of 35 sites. For more detailed information on data collection through passive acoustic monitoring, see Bueno et al. (2020).

### 3.2. Compound taxonomic diversity of soniferous taxa

To assess the relationship between the soundscape diversity metrics and the taxonomic richness of sound-producing organisms in the study area, we generated an index capturing the compound species richness of three major tropical forest soniferous taxa: (i) anurans; (ii) birds; and (iii) primates. We characterised the taxonomic richness of anurans using 62 1-min recordings per plot, considering 1 min every 10 mins between 17:00h – 22:01h, a period within which anurans are known to be most sonically active, during sampling days 2 and 4 out of 5 days. Anuran species data were obtained through visual and aural identification of anuran sounds by an expert observer (GSM) using the RFCx Arbimon II Visualizer Tool (https://arbimon.rfcx.org/). Species identity was cross-verified by a second expert observer (ILK) to ensure accuracy. For bird species data, we used for each plot a combination of aural identification by a trained expert (MC-C) and a Pattern Matching algorithm available in RFCx Arbimon to automatically detect species in the sound files described in section 3.1. For the taxonomic richness of primates, we compiled data from a previously published study on vertebrate diversity at the Balbina Hydroelectric Dam (Benchimol and Peres 2015, 2021). We consequently retained only sites for which taxonomic richness data of all three groups were available (Table S3; see S.6 for detailed information).

For the sampling sites containing multiple plots, the cumulative taxonomic richness across plots was calculated for all taxonomic groups. A total of 34 anuran species, 71 bird species and 7 primate species were found across the 35 sites. Finally, for each island and continuous forest site, the richness values for anurans, birds and primates were summed to obtain the compound richness of soniferous species. We acknowledge that this compound richness does not include insects, which represent a dominant acoustic group in tropical forests (Aide et al. 2017). Still, we deem the combined acoustic activity of these three taxonomic groups to be sufficiently strong to influence the rain forest soundscape, and therefore detectable by our soundscape diversity metrics.

### 3.3. Soundscape diversity data

To derive the soundscape diversity metrics, the set of recordings described in section 3.1 was used (see Table S3 for details). Per plot, we quantified the presence of sound in the acoustic trait space by computing the CVR-index for each 1-min file using a 256-frame window length (see S.2. for more information on window length choice) and 44,100 Hz sampling rate. This measure quantifies the fraction of cells in each noise-reduced frequency bin that exceed a 3 dB amplitude value. Next, we concatenated the CVR files chronologically per plot. Then, we determined a binarisation threshold for each plot using the ‘IsoData’ binarisation algorithm in the ‘autothresholdr’ R-package (v1.3.11; Landini et al. 2017; see S.3 for more details). Using this site-specific threshold, we binarised the CVR-values to obtain a detection (1) / non-detection (0) variable for each OSU (see S.3. for a detailed breakdown of thresholding methods). After, we separated the binarised spectral index files into 24h samples (288 files using a 1/5 min sampling regime) of the soundscape. Finally, an OSU-by-sample incidence matrix for each plot was obtained. For sampling sites containing multiple plots, the obtained OSU-by-sample incidence matrices were grouped across plots.

As highlighted in Metcalf et al. (2020), *a priori* knowledge of acoustic space usage can be used to subset the acoustic trait space to those time-frequency coordinates used by the soniferous groups of interest. Doing so can help avoid signal masking, thus increasing the sensitivity of soundscape metrics to taxonomic richness. As such, knowing that the upper frequency range of the sonically active groups used in our study is 14,384 Hz (*Osteocephalus buckleyi*), we subset the frequency domain below 14,125 Hz. Moreover, as the sampling duration is unequal between plots in the study, and we wish to retain the maximal amount of information without discarding data unnecessarily, we used sample size-based interpolation/extrapolation curves to equalise the sampling effort among plots (see S.1. for more information on rarefaction). We are interested in characterising the exact relationships in the soundscape diversity metrics between plots, so we can extrapolate to double the minimal sample size at most (Chao and Jost 2012). To achieve this, we use the R-package *‘iNEXT’* (Hsieh et al. 2016) to calculate soundscape richness (*^0^D*) and evenness (*^2^D* / *^0^D*) at a sampling effort of 8 days (twice the minimal sampling duration). Finally, by computing the Pearson correlation coefficient and R^2^-value for a simple linear regression model, we investigated the relationship between the soundscape richness and evenness, and the independently sampled compound richness of soniferous species. To assess the robustness of the workflow, we provide additional analyses on the effect of sampling regime (S.1.2.2) and window length (S.2) on the relationship between soundscape richness and taxonomic richness in the supplementary materials.

## 4. Results

### 4.1. Properties of soundscape diversity metrics

The ensemble of all three indices has strictly positive values, can be constrained between 0-1, and is theoretically independent of the taxonomic richness (S.5.1). Moreover, the monotonicity criterion holds true for the soundscape richness and diversity metrics, but not for the soundscape evenness (S.5.2). The soundscape richness and evenness are independent from one another and describe unique aspects of the soundscape diversity (S.5.3). Conversely, the soundscape diversity at *q* = 2 displays a positive relationship with both the soundscape richness and evenness, and thus does not conform with the independence criterion. Additionally, unlike some commonly used biodiversity indices such as the Shannon-Wiener and Simpson biodiversity index, our metrics scale linearly with the underlying diversity of the system - a theorem known as the replication principle (S.5.4). Finally, the same analytical workflow can be used to quantify the soundscape diversity at multiple scales or hierarchical levels, decomposing the regional metacommunity diversity (γ-diversity) into its local diversity (α-diversity) and community turnover (β-diversity) components using a simple multiplicative relationship (S.4; S.5.6).

### 4.2. Relationship between soundscape metrics and taxonomic richness

As indicated by the Pearson correlation coefficient and associated R^2^- and p-values, the correlation between the soundscape richness and richness of soniferous species is positive and strong (*r* = 0.84; R^2^ = 0.71; p < 0.001; Fig. 4A-1; Table S4). This correlation holds even for lower intensity sampling regimes (S.1.2.2), with values staying high (r > 0.8) for all tested sampling intensities. Moreover, we find that the choice of window length has a negligible impact on the observed correlation between both metrics (*r* > 0.83 for all window lengths; see S.2). Additionally, based on visual inspection of the acoustic trait space, islands containing a lower richness of soniferous species (Fig. 4A-2) appear to have more empty and less complex trait spaces than species-rich islands (Fig. 4A-3). The trait space of low richness islands has impoverished daytime soundscapes and lacks many of the sounds exceeding 5,000 Hz that are present at taxonomically rich islands. For the soundscape evenness, the correlation with the richness of soniferous species is weakly positive (*r* = 0.35; R^2^ = 0.12; p < 0.041: Fig. 4B-1). A visual of the acoustic trait space reveals that, for low evenness sites (Fig. 4B-2), low abundance sounds are much more prevalent compared to sites with a high soundscape evenness (Fig. 4B-3).

**Figure 4:**
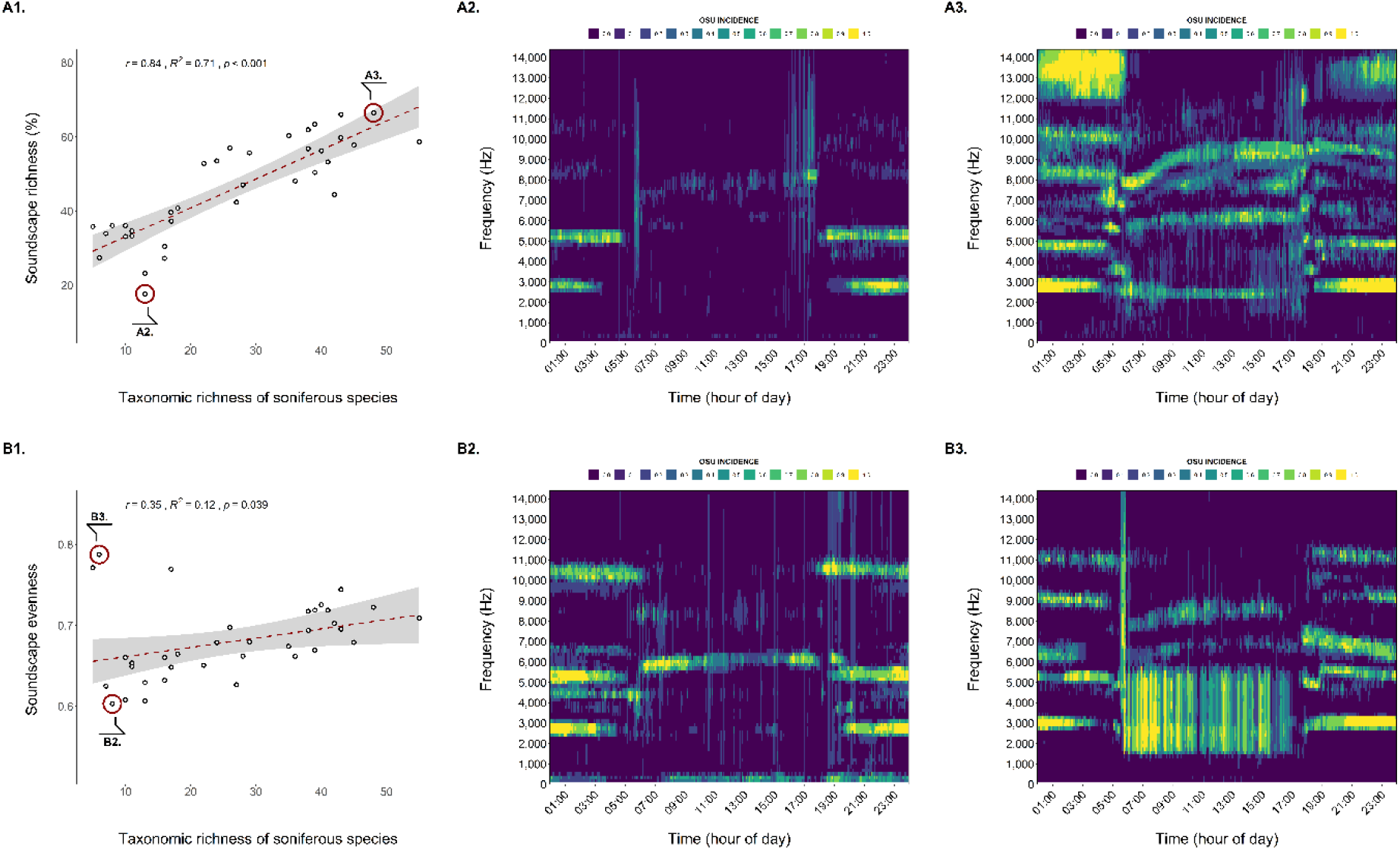
**A.** Relationship between the soundscape richness and the richness of soniferous species (**A1**) with a visual representation of the 24-hour acoustic trait space for low-richness (**A2**) and high-richness (**A3**) soundscapes. The Pearson correlation coefficient and associated R^2^- and p-values indicate a strong positive relationship (r = 0.84) between the soundscape richness and species richness of sound-producing vertebrates. **B.** The relationship between the soundscape evenness and the richness of soniferous species (**B1**) with a visual representation of low-evenness (**B2**) and high-evenness (**B3**) soundscapes. The Pearson correlation coefficient and associated R^2^- and p-values indicate a weak positive correlation (r = 0.35) between the soundscape evenness and taxonomic richness.

## 5. Discussion

In this manuscript, we applied the principles of functional diversity research to the realm of soundscape ecology, proposing a novel workflow to study patterns in the 24-hour acoustic trait space usage at a landscape scale. Moreover, we assessed the ability of these soundscape diversity metrics to capture patterns in the richness of the soniferous community. To do so, we used a spectral acoustic index to capture the functional traits of sounds without the need to isolate and identify individual species calls. Instead, we group sounds by their shared spectro-temporal amplitude features in functional trait space, thereby creating a novel unit of soundscape diversity measurement – the Operation Sound Unit (OSU). Next, we quantified the presence and abundance of these OSUs throughout the recording period and used the framework of Hill numbers to decompose their diversity into measures of soundscape richness, diversity, and evenness.

### 5.1. Advantages of the workflow

First, we show that the proposed soundscape diversity metrics abide by a set of fundamental criteria for functional diversity indices (Ricotta 2005; Villéger et al. 2008; Mouchet et al. 2010) and behave in an ecologically intuitive manner. Second, breaking down the soundscape diversity into its various facets (richness, evenness and diversity) and assessing how they behave along a gradient in the taxonomic richness of the acoustic community can shed light on patterns of acoustic niche usage and the processes influencing them. In soundscape ecology, two prevailing hypotheses are believed to influence acoustic community assembly and niche usage: (i) the Acoustic Adaptation Hypothesis; (ii) the Acoustic Niche Hypothesis (Pijanowski et al. 2011a). The former posits that species’ acoustic functional traits (e.g., signal frequency, amplitude, timing, and duration) are more similar than expected by chance as the environment filters for traits that maximise effective sound propagation and minimise attenuation in the physical habitat (Mullet et al. 2017). The latter states that the acoustic trait space is a core ecological resource and sonically sympatric species are thought to partition their acoustic niche in the time and frequency domains to avoid spectro-temporal overlap in their sounds, which would lead to inefficient communication (Krause 1993; Garcia-Rutledge and Narins 2001). Moreover, the Acoustic

Niche Hypothesis implies that evolutionarily archaic and undisturbed ecosystems have acquired an evolutionary balance between all sounds in the landscape, resulting in soundscapes with high spectro-temporal complexity and signal diversity, and minimal overlap (Krause 1993; Pijanowski et al. 2011a; Eldridge et al. 2016). Conversely, disturbed systems in which *‘acoustically optimised’* species have been lost from the habitat are then characterised by an unbalanced equilibrium, showing readily detectable gaps in the soundscape.

The soundscape richness metric described herein quantifies the amount of acoustic niche space that is occupied by OSUs independent of their relative abundances. In our case study, we found a strong positive correlation (*r* = 0.84; R^2^ = 0.71; p < 0.001) between the soundscape richness and the species richness of the soniferous community. As demonstrated, the soundscape richness is theoretically independent of the species richness, so the observed relationship likely arises through processes of species assembly. Following the Acoustic Adaptation Hypothesis, we would expect the richness of OSUs, driven by the richness of acoustic functional trait values, to be mostly insensitive to the richness of soniferous species. As such, considering the strong positive relationship between the soundscape richness and the richness of soniferous species, it is likely that the acoustic community in our study is structured by competition for acoustic niche space. As the taxonomic richness gradient in our study area originated from a disturbance event, it is plausible that the observed correlations between the soundscape richness and taxonomic richness stem from the loss of species occupying unique acoustic niches in the acoustic trait space, resulting in a lower niche saturation at lower taxonomic richness.

The soundscape evenness metric captures the degree to which the relative abundances of OSUs are distributed in niche space. Thus, it quantifies how effectively the available acoustic resources are used at a landscape scale and sheds light on patterns of dominance and rarity. In the case study, the soundscape evenness displays a weak positive correlation (*r* = 0.35; R^2^ = 0.12: p < 0.05) with the richness of soniferous species. This suggests that changes in the richness of soniferous species bring about associated changes in the relative abundance distribution of sounds in acoustic trait space. We posit that the observed correlation between the soundscape evenness and richness of soniferous species could reflect an unbalanced equilibrium, in which the acoustic community consists of a few acoustically dominant and many rare sound producing species (Krause 1993). As such, it appears that disturbed species-poor acoustic communities use the acoustic niche resource less effectively (Mason et al. 2005). Clearly, the combination of both richness and evenness metrics can provide unique insights into acoustic niche usage. Yet many of the existing soundscape diversity metrics focus solely on the presence of sound in a short duration recording without accounting for the distribution of sound abundance throughout the recording period, thus potentially overlooking the evenness component of soundscape diversity.

Third, we see potential to use the workflow described herein as a robust and cost-effective method to track biodiversity changes at large spatial and temporal scales or in systems where the knowledge of the resident biological community is incomplete, particularly where labour-intensive and time-consuming traditional biodiversity monitoring methods may be unfeasible. As we demonstrate a strong positive correlation between the soundscape richness and an independent estimate of ground-truthed taxonomic diversity of soniferous species, this metric can be used as a proxy to infer taxonomic diversity patterns. In this way, it can be used as an early warning system, alerting researchers when declines in soundscape diversity exceed natural fluctuations (Pijanowski et al. 2011a; Krause and Farina 2016). Compared to analogous metrics described in the literature which capture acoustic niche usage, the soundscape richness metric performs well as a biodiversity proxy. For instance, in Burivalova et al. (2019), the soundscape saturation, which calculates the saturation of acoustic niche space for 1-min duration sound files, achieves a correlation of r = 0.55 and R^2^ = 0.31 with the taxonomic richness of vertebrates identified in the same sound file. Moreover, the soundscape richness metric described herein reaches equivalent correlations (Spearman’s ρ ≈ 0.85) to the Acoustic Space Use metric described in Aide et al. (2017), yet the latter study had a relatively small sample size (8 plots).

Furthermore, even when a correlation between soundscape diversity metrics and taxonomic diversity is absent, the workflow can still inform ecological research. It allows us to measure where and when in the acoustic trait space the occurrence and relative abundance distribution of sound is changing across space, time, or hierarchical levels (e.g., local, regional, or global) without requiring a link to the taxonomic identity of OSUs. For instance, in our case study, we visually compared differences in acoustic trait space usage between two extremes of the richness gradient and found that low richness sites had an impoverished daytime soundscape and lacked sounds over 5,000 Hz. Moreover, the low evenness soundscape had a higher proportion of rare OSUs, suggesting the acoustic niche resource was used less effectively. This knowledge could be used in synergy with the aural investigation of sound files, directing species identification to the regions of acoustic trait space where a change in OSU presence and incidence has been observed. This, in turn, can direct field-based studies using traditional monitoring methodologies towards the taxonomic groups for which change was discerned.

Finally, we demonstrated that the workflow we propose here is robust, picking up on an ecological gradient in an acoustically complex tropical rain forest setting. We were not interested in non-biological or biological low-amplitude sounds which emanate from long distances. As such, we removed these sounds from the data by using a 3 dB amplitude threshold to compute the CVR-index. Additionally, we had no interest in transient/short-duration high-amplitude sounds in otherwise silent frequency bins, as their impact on the acoustic space is negligible and potentially of non-biological origin. We therefore converted CVR values to incidence data by only considering an OSU as detected (1) when a certain proportion of cells in the frequency bin exceeds a 3 dB threshold. We argue this will reduce potential differences between sites and eliminate the non-focal transient and low amplitude sounds from the data. Although these steps do not remove persistent non-focal high amplitude sounds such as rain showers, thunder, or wind, from the data, we still find strong positive correlations with taxonomic richness without actively removing geophonic events from the recordings.

Moreover, we found that the choice of window length for spectral index computation has a minimal effect on the observed correlation between the soundscape richness and richness of soniferous species. We also discovered that at equal sampling duration, this correlation remains equally strong (*r* = 0.84) for medium intensity sampling regimes (1/10 min; 1/15 min; 1/20 min), and only slightly diminished (*r* ≥ 0.82) for low-intensity sampling regimes (1/30 min; 1/60 min). At last, in our study, the soundscape richness ranking among sites stabilized after 8-10 sampling days. Knowing that our most intense sampling regime was 1 min / 5 min, this corresponds to 38.4 to 48 hours of recordings per site. This number of recording hours associated with the sampling effort required to capture a meaningful soundscape reliably is considerably lower than other efforts previously reported in the literature (*e.g.* 120 hours in Bradfer-Lawrence et al. 2019). If the sampling design is determined by the available storage on the acoustic sensor’s memory card, this is good news, as cards with a lower capacity could be used or the excess storage space could be allocated to sampling the soundscape at higher sampling rates rather than for more days or higher intensity regimes.

### 5.2. Avenues of future research

At present, the soundscape diversity metrics outlined herein treat all OSUs as equally similar. In reality, OSUs are not independent elements, but rather correlated units in acoustic trait space. As such, future work on soundscape diversity metrics should incorporate the difference in acoustic trait values (time-frequency coordinates) of a particular OSU from all other OSUs in the acoustic space (Scheiner et al. 2017). Incorporating the distinctiveness of OSUs in acoustic trait space - which we term soundscape dispersion - would allow us to further quantify the degree to which acoustic trait space is partitioned, providing further insights into acoustic niche differentiation and resource competition (Mason et al. 2005). For instance, if acoustic communities are structured by competition for acoustic space, we might expect overdispersion in acoustic trait space compared to the same number of OSUs drawn randomly from the regional OSU pool. Conversely, when the dispersion of OSUs in acoustic space is lower than expected compared to the randomly drawn OSU pool, environmental filtering is likely to be an acting process (Scheiner et al. 2017).

In this work, we opted for an incidence-based approach to attribute an importance value to OSUs. Yet, the use of threshold values to convert continuous variables to detection/non-detection data has been critiqued in the literature (Lawson et al. 2014), as it results in information loss and complicates comparisons between different sites/studies for which different optimal threshold values may apply. Still, we posit this approach can be appropriate for soundscape data. Although acoustic indices are known to capture animal activity, there is an ongoing debate about their ability to capture patterns of abundance (Boelman et al. 2007; Bradfer-Lawrence et al. 2020). Moreover, acoustic indices can be sensitive to confounding environmental factors (Gasc et al. 2015). For instance, CVR-index values may respond to abiotic sounds, such as geophony and anthrophony, which are considered confounding factors if the aim is to capture biophonic sounds. Additionally, the index values can also be susceptible to the relative amplitude of songs in recordings, which in turn are shaped by the properties of the surrounding vegetation, the distance of the sound-emitting animal to the sensor, inherent biological differences between species, and meteorological conditions (Bradfer-Lawrence et al. 2020). Some of these external factors can vary greatly between sites, which could challenge the trustworthiness of raw observed differences in index values. We argue that converting raw CVR-index values to binary detection/non-detection data will reduce potential differences between sites and eliminate the non-focal transient and low amplitude sounds from the data.

Nonetheless, choosing a threshold value that is valid in all ecological contexts and accurately removes non-target sounds while retaining enough information to capture patterns in niche usage represents a challenge. Deriving a unique threshold value for each study system by validating the ability of the soundscape diversity metrics to capture a species richness gradient is not a feasible approach, as this taxonomic data will not always be available. As such, in S.3, we compare various thresholding methods and assess their utility for our workflow. We find that the approach described in Burivalova et al. (2018), for which the chosen threshold yields the most normal distribution of obtained soundscape metric, does not yield the strongest correlation values with species richness. Moreover, although we find that using a constant threshold value of 0.09 across all sites works well for our specific case study, it is likely that this threshold value will be different for other ecosystems, seasons, or levels of non-target sound. Instead, we recommend that future studies derive incidence data by using context-aware binarisation algorithms (see S.3). These algorithms produce a unique binarisation threshold per site by considering the distribution of CVR-values in the acoustic trait space, which in turn is influenced by the soniferous community and sound transmission characteristics of the habitat. We find that the ‘IsoData’ binarisation algorithm works best for our data, but further research in a wider variety of habitats is needed to confirm that this algorithm performs most consistently.

Finally, in this work, we used the CVR-index to capture the general amplitude features inherent to all sounds produced in the landscape. We posit that, by exploring other spectral indices or combinations of indices, each capturing the structure and distribution of acoustic energy that is characteristic of a specific taxonomic group of interest, the sensitivity to these groups might be increased. For instance, cicada choruses are characterised by loud and long-duration stridulations, usually restricted to narrow frequency bands. Previous work suggests these features can be captured by a set of spectral indices: (i) low spectral entropy; (ii) high background noise; (iii) high spectral density (Ferroudj et al. 2014; Towsey et al. 2014; Brown et al. 2019). As such, we argue that, if the soundscape diversity of cicada chorusing is of interest to the study, these three spectral indices could be combined into a compound cicada index, which could then be used to decompose the diversity of cicada choruses in the 24-hour acoustic trait space. Similarly, we hypothesise that other spectral indices or combinations of indices could be used to target specific groups of interest by exploiting the ability of these indices to capture specific amplitude features in the time-frequency domain.

## 6. Conclusion

In this work, we provide a workflow for the quantification of soundscape diversity which builds on functional ecology and uses the analytical framework of Hill numbers to generate a robust set of soundscape diversity metrics. By broadening the temporal scope of soundscape diversity quantification to the 24h period in which all species can produce sound and considering the spectral and temporal traits of sound simultaneously, these soundscape diversity metrics can yield novel insights into acoustic trait space usage at multiple spatio-temporal scales and act as a proxy for the richness of soniferous species in an acoustically complex environment.

## 7. Author contributions

TL designed the workflow, coded the accompanying package, analysed the case study and supplementary material data, and wrote the manuscript. ASB, TH and CAP made significant contributions in the study conception and design, and manuscript revision. ASB and CAP managed the design and collection of acoustic data. GSM and ILK performed identification of anuran species in sound files and revised the final manuscript. MC-C performed the aural identification of bird species in sound files and revised the final manuscript.

## 8. Acknowledgements

We thank the two anonymous reviewers for their constructive comments which considerably improved the manuscript quality. For their assistance in data collection, we are very grateful to Evanir Damasceno, Tatiane Abreu, and Carla Fonseca. We are thankful to the staff at Reserva Biológica do Uatumã for logistical support. Data collection was funded by the Rufford Foundation (grant 17715-1), Reserva Biológica do Uatumã (ICMBio), the University of East Anglia and a NERC/UK grant (NE/J01401X/1) awarded to CAP. ASB was funded by a PhD studentship (grant 200463/2014-4) from Conselho Nacional de Desenvolvimento Científico e Tecnológico (CNPq) – Brazil, which also awarded a productivity grant to ILK (309415/2016-0). TL is supported by a PhD studentship from the Norwegian University of Life Sciences.

## Publication bibliography

Agostino, Patricia V.; Lusk, Nicholas A.; Meck, Warren H.; Golombek, Diego A.; Peryer, Guy (2020): Daily and seasonal fluctuation in Tawny Owl vocalization timing. In PloS one 15 (4), e0231591. DOI: 10.1371/journal.pone.0231591.

Aide, T.; Hernández-Serna, Andres; Campos-Cerqueira, Marconi; Acevedo-Charry, Orlando; Deichmann, Jessica (2017): Species Richness (of Insects) Drives the Use of Acoustic Space in the Tropics. In Remote Sensing 9 (11), p. 1096. DOI: 10.3390/rs9111096.

Benchimol, Maíra; Peres, Carlos A. (2015): Widespread Forest Vertebrate Extinctions Induced by a Mega Hydroelectric Dam in Lowland Amazonia. In PLOS ONE 10 (7), e0129818. DOI: 10.1371/journal.pone.0129818.

Benchimol, Maíra; Peres, Carlos A. (2021): Determinants of population persistence and abundance of terrestrial and arboreal vertebrates stranded in tropical forest land-bridge islands. In Conservation Biology 35 (3), pp. 870–883. DOI: 10.1111/cobi.13619.

Boelman, Natalie T.; Asner, Gregory P.; Hart, Patrick J.; Martin, Roberta E. (2007): Multi-trophic invasion resistance in Hawaii: bioacoustics, field surveys, and airborne remote sensing. In Ecological applications: a publication of the Ecological Society of America 17 (8), pp. 2137–2144. DOI: 10.1890/07-0004.1.

Bradfer-Lawrence, Tom; Bunnefeld, Nils; Gardner, Nick; Willis, Stephen G.; Dent, Daisy H. (2020): Rapid assessment of avian species richness and abundance using acoustic indices. In Ecological Indicators 115, p. 106400. DOI: 10.1016/j.ecolind.2020.106400.

Bradfer-Lawrence, Tom; Gardner, Nick; Bunnefeld, Lynsey; Bunnefeld, Nils; Willis, Stephen G.; Dent, Daisy H. (2019): Guidelines for the use of acoustic indices in environmental research. In Methods Ecol Evol 10 (10), pp. 1796–1807. DOI: 10.1111/2041-210X.13254.

Brown, Alexander; Garg, Saurabh; Montgomery, James (2019): Automatic rain and cicada chorus filtering of bird acoustic data. In Applied Soft Computing 81, p. 105501. DOI: 10.1016/j.asoc.2019.105501.

Bueno, Anderson Saldanha; Masseli, Gabriel S.; Kaefer, Igor L.; Peres, Carlos A. (2020): Sampling design may obscure species-area relationships in landscape-scale field studies. In Ecography 43 (1), pp. 107–118. DOI: 10.1111/ecog.04568.

Burivalova, Zuzana; Purnomo; Wahyudi, Bambang; Boucher, Timothy M.; Ellis, Peter; Truskinger, Anthony et al. (2019): Using soundscapes to investigate homogenization of tropical forest diversity in selectively logged forests. In J Appl Ecol 56 (11), pp. 2493–2504. DOI: 10.1111/1365-2664.13481.

Burivalova, Zuzana; Towsey, Michael; Boucher, Tim; Truskinger, Anthony; Apelis, Cosmas; Roe, Paul; Game, Edward T. (2018): Using soundscapes to detect variable degrees of human influence on tropical forests in Papua New Guinea. In Conservation biology: the journal of the Society for Conservation Biology 32 (1), pp. 205–215. DOI: 10.1111/cobi.12968.

Buxton, Rachel T.; McKenna, Megan F.; Clapp, Mary; Meyer, Erik; Stabenau, Erik; Angeloni, Lisa M. et al. (2018): Efficacy of extracting indices from large-scale acoustic recordings to monitor biodiversity. In Conservation biology: the journal of the Society for Conservation Biology 32 (5), pp. 1174–1184. DOI: 10.1111/cobi.13119.

Chao, Anne; Chiu, Chun-Huo; Hsieh, T. C. (2012): Proposing a resolution to debates on diversity partitioning. In Ecology 93 (9), pp. 2037–2051. DOI: 10.1890/11-1817.1.

Chao, Anne; Chiu, Chun-Huo; Jost, Lou (2014): Unifying Species Diversity, Phylogenetic Diversity, Functional Diversity, and Related Similarity and Differentiation Measures Through Hill Numbers. In Annu. Rev. Ecol. Evol. Syst. 45 (1), pp. 297–324. DOI: 10.1146/annurev-ecolsys-120213-091540.

Chao, Anne; Chiu, Chun-Huo; Villéger, Sébastien; Sun, I-Fang; Thorn, Simon; Lin, Yi-Ching et al. (2019): An attribute-diversity approach to functional diversity, functional beta diversity, and related (dis)similarity measures. In Ecol Monogr 89 (2), e01343. DOI: 10.1002/ecm.1343.

Chao, Anne; Jost, Lou (2012): Coverage-based rarefaction and extrapolation: standardizing samples by completeness rather than size. In Ecology 93 (12), pp. 2533–2547. DOI: 10.1890/11-1952.1.

Chiu, Chun-Huo; Chao, Anne (2014): Distance-based functional diversity measures and their decomposition: a framework based on Hill numbers. In PloS one 9 (7), e100014. DOI: 10.1371/journal.pone.0100014.

Da Silva, Crhistiane Andressa; Pontes, André Luiz Bezerra de; Cavalcante, Jeferson de Souza; Azevedo, Carolina Virgínia Macedo de (2014): Conspecific vocalisations modulate the circadian activity rhythm of marmosets. In Biological Rhythm Research 45 (6), pp. 941–954. DOI: 10.1080/09291016.2014.939441.

Darras, Kevin; Pütz, Peter; Fahrurrozi; Rembold, Katja; Tscharntke, Teja (2016): Measuring sound detection spaces for acoustic animal sampling and monitoring. In Biological Conservation 201, pp. 29–37. DOI: 10.1016/j.biocon.2016.06.021.

Darwin, Charles (1872): The expression of the emotions in man and animals. In London, UK: John Marry.

Depraetere, Marion; Pavoine, Sandrine; Jiguet, Fréderic; Gasc, Amandine; Duvail, Stéphanie; Sueur, Jérôme (2012): Monitoring animal diversity using acoustic indices: Implementation in a temperate woodland. In Ecological Indicators 13 (1), pp. 46–54. DOI: 10.1016/j.ecolind.2011.05.006.

Eldridge, Alice; Casey, Michael; Moscoso, Paola; Peck, Mika (2016): A new method for ecoacoustics? Toward the extraction and evaluation of ecologically-meaningful soundscape components using sparse coding methods. In PeerJ 4, e2108. DOI: 10.7717/peerj.2108.

Farina, Almo (2013): Soundscape ecology: principles, patterns, methods and applications: Springer.

Farina, Almo; Ceraulo, Maria; Bobryk, Christopher; Pieretti, Nadia; Quinci, Enza; Lattanzi, Emanuele (2015): Spatial and temporal variation of bird dawn chorus and successive acoustic morning activity in a Mediterranean landscape. In Bioacoustics 24 (3), pp. 269–288. DOI: 10.1080/09524622.2015.1070282.

Farina, Almo; James, Philip (2016): The acoustic communities: Definition, description and ecological role. In Biosystems 147, pp. 11–20. DOI: 10.1016/j.biosystems.2016.05.011.

Ferroudj, Meriem; Truskinger, Anthony; Towsey, Michael; Zhang, Liang; Zhang, Jinglan; Roe, Paul (2014): Detection of Rain in Acoustic Recordings of the Environment. In Duc-Nghia Pham, Seong-Bae Park (Eds.): PRICAI 2014: Trends in Artificial Intelligence. Cham, 2014. Cham: Springer International Publishing, pp. 104–116.

Fuller, Susan; Axel, Anne C.; Tucker, David; Gage, Stuart H. (2015): Connecting soundscape to landscape: Which acoustic index best describes landscape configuration? In Ecological Indicators 58, pp. 207–215. DOI: 10.1016/j.ecolind.2015.05.057.

Garcia-Rutledge, Elizabeth J; Narins, Peter M. (2001): Shared Acoustic Resources in an Old World Frog Community. In Herpetologica 57 (1), pp. 104–116. Available online at http://www.jstor.org/stable/3893144.

Gasc, A.; Pavoine, S.; Lellouch, L.; Grandcolas, P.; Sueur, J. (2015): Acoustic indices for biodiversity assessments: Analyses of bias based on simulated bird assemblages and recommendations for field surveys. In Biological Conservation 191, pp. 306–312. DOI: 10.1016/j.biocon.2015.06.018.

Gasc, A.; Sueur, J.; Jiguet, F.; Devictor, V.; Grandcolas, P.; Burrow, C. et al. (2013): Assessing biodiversity with sound: Do acoustic diversity indices reflect phylogenetic and functional diversities of bird communities? In Ecological Indicators 25, pp. 279–287. DOI: 10.1016/j.ecolind.2012.10.009.

Gibb, Rory; Browning, Ella; Glover-Kapfer, Paul; Jones, Kate E. (2019): Emerging opportunities and challenges for passive acoustics in ecological assessment and monitoring. In Methods Ecol Evol 10 (2), pp. 169–185. DOI: 10.1111/2041-210X.13101.

Hill, M. O. (1973): Diversity and Evenness: A Unifying Notation and Its Consequences. In Ecology 54 (2), pp. 427–432. DOI: 10.2307/1934352.

Hsieh, T. C.; Ma, K. H.; Chao, Anne (2016): iNEXT: an R package for rarefaction and extrapolation of species diversity (H ill numbers). In Methods Ecol Evol 7 (12), pp. 1451–1456. DOI: 10.1111/2041-210X.12613.

Jianguo, C. U.I.; Xiaoyan, SONG; Guangzhan, FANG; Fei, X. U.; Steven E., BRAUTH; Yezhong, TANG (2011): Circadian Rhythm of Calling Behavior in the Emei Music Frog *(Babina daunchina)* is Associated with Habitat Temperature and Relative Humidity. In Asian Herpetological Research 2 (3), pp. 149–154. DOI: 10.3724/SP.J.1245.2011.00149.

Jost, Lou (2006): Entropy and diversity. In Oikos 113 (2), pp. 363–375. DOI: 10.1111/j.2006.0030-1299.14714.x.

Jost, Lou (2007): Partitioning diversity into independent alpha and beta components. In Ecology 88 (10), pp. 2427–2439. DOI: 10.1890/06-1736.1.

Jost, Lou (2010): The Relation between Evenness and Diversity. In Diversity 2 (2), pp. 207–232. DOI: 10.3390/d2020207.

Krause, Bernard L. (1993): The niche hypothesis: a virtual symphony of animal sounds, the origins of musical expression and the health of habitats. In The Soundscape Newsletter 6, pp. 6–10.

Krause, Bernie (1987): Bioacoustics, habitat ambience in ecological balance. In Whole Earth Review 57 (Winter).

Krause, Bernie; Farina, Almo (2016): Using ecoacoustic methods to survey the impacts of climate change on biodiversity. In Biological Conservation 195, pp. 245–254. DOI: 10.1016/j.biocon.2016.01.013.

Landini, G.; Randell, D. A.; Fouad, S.; Galton, A. (2017): Automatic thresholding from the gradients of region boundaries. In Journal of microscopy 265 (2), pp. 185–195. DOI: 10.1111/jmi.12474.

Lawson, Callum R.; Hodgson, Jenny A.; Wilson, Robert J.; Richards, Shane A. (2014): Prevalence, thresholds and the performance of presence-absence models. In Methods Ecol Evol 5 (1), pp. 54–64. DOI: 10.1111/2041-210X.12123.

Mason, Norman W. H.; Mouillot, David; Lee, William G.; Wilson, J. Bastow (2005): Functional richness, functional evenness and functional divergence: the primary components of functional diversity. In Oikos 111 (1), pp. 112–118. DOI: 10.1111/j.0030-1299.2005.13886.x.

Metcalf, Oliver C.; Barlow, Jos; Devenish, Christian; Marsden, Stuart; Berenguer, Erika; Lees, Alexander C. (2020): Acoustic indices perform better when applied at ecologically meaningful time and frequency scales. In Methods Ecol Evol. DOI: 10.1111/2041-210X.13521.

Mouchet, Maud A.; Villéger, Sébastien; Mason, Norman W. H.; Mouillot, David (2010): Functional diversity measures: an overview of their redundancy and their ability to discriminate community assembly rules. In Functional Ecology 24 (4), pp. 867–876. DOI: 10.1111/j.1365-2435.2010.01695.x.

Mullet, Timothy C.; Farina, Almo; Gage, Stuart H. (2017): The Acoustic Habitat Hypothesis: An Ecoacoustics Perspective on Species Habitat Selection. In Biosemiotics 10 (3), pp. 319–336. DOI: 10.1007/s12304-017-9288-5.

Pavoine, S.; Bonsall, M. B. (2011): Measuring biodiversity to explain community assembly: a unified approach. In Biological reviews of the Cambridge Philosophical Society 86 (4), pp. 792–812. DOI: 10.1111/j.1469-185X.2010.00171.x.

Pieretti, N.; Farina, A.; Morri, D. (2011): A new methodology to infer the singing activity of an avian community: The Acoustic Complexity Index (ACI). In Ecological Indicators 11 (3), pp. 868–873. DOI: 10.1016/j.ecolind.2010.11.005.

Pijanowski, Bryan C.; Farina, Almo; Gage, Stuart H.; Dumyahn, Sarah L.; Krause, Bernie L. (2011a): What is soundscape ecology? An introduction and overview of an emerging new science. In Landscape Ecol 26 (9), pp. 1213–1232. DOI: 10.1007/s10980-011-9600-8.

Pijanowski, Bryan C.; Villanueva-Rivera, Luis J.; Dumyahn, Sarah L.; Farina, Almo; Krause, Bernie L.; Napoletano, Brian M. et al. (2011b): Soundscape Ecology: The Science of Sound in the Landscape. In BioScience 61 (3), pp. 203–216. DOI: 10.1525/bio.2011.61.3.6.

R Core Team (2020): R: A Language and Environment for Statistical Computing. Vienna, Austria. Available online at https://www.r-project.org/.

Ricotta, Carlo (2005): A note on functional diversity measures. In Basic and Applied Ecology 6 (5), pp. 479–486. DOI: 10.1016/j.baae.2005.02.008.

Rodriguez, Alexandra; Gasc, Amandine; Pavoine, Sandrine; Grandcolas, Philippe; Gaucher, Philippe; Sueur, Jérôme (2014): Temporal and spatial variability of animal sound within a neotropical forest. In Ecological Informatics 21, pp. 133–143. DOI: 10.1016/j.ecoinf.2013.12.006.

Scheiner, Samuel M.; Kosman, Evsey; Presley, Steven J.; Willig, Michael R. (2017): Decomposing functional diversity. In Methods Ecol Evol 8 (7), pp. 809–820. DOI: 10.1111/2041-210X.12696.

Seyfarth, Robert M.; Cheney, Dorothy L. (2003): Meaning and emotion in animal vocalizations. In Annals of the New York Academy of Sciences 1000, pp. 32–55. DOI: 10.1196/annals.1280.004.

Sokal, Robert R.; Sneath, Peter H. A. (1963): Principles of numerical taxonomy.

Sugai, Larissa Sayuri Moreira; Silva, Thiago Sanna Freire; Ribeiro, José Wagner; Llusia, Diego (2019): Terrestrial Passive Acoustic Monitoring: Review and Perspectives. In BioScience 69 (1), pp. 15–25. DOI: 10.1093/biosci/biy147.

Toledo, Luís Felipe; Tipp, Cheryl; Márquez, Rafael (2015): The value of audiovisual archives. In Science (New York, N.Y.) 347 (6221), p. 484. DOI: 10.1126/science.347.6221.484-b.

Towsey, Michael (2017): The calculation of acoustic indices derived from long-duration recordings of the natural environment. Available online at https://eprints.qut.edu.au/110634/.

Towsey, Michael; Truskinger, Anthony; Cottman-Fields, Mark; Roe, Paul (2018): Ecoacoustics Audio Analysis Software V18.03.0.41: Zenodo. Available online at https://ap.qut.ecoacoustics.info/tutorials/01-usingap/practical?tabs=windows;

Towsey, Michael; Wimmer, Jason; Williamson, Ian; Roe, Paul (2014): The use of acoustic indices to determine avian species richness in audio-recordings of the environment. In Ecological Informatics 21, pp. 110–119. DOI: 10.1016/j.ecoinf.2013.11.007.

Truskinger, Anthony; Towsey, Michael (2019): Why do we analyse data in 1-min chunks? Available online at https://research.ecosounds.org/2019/08/09/analyzing-data-in-one-minute-chunks.html, checked on 9/9/2019.

Tucker, David; Gage, Stuart H.; Williamson, Ian; Fuller, Susan (2014): Linking ecological condition and the soundscape in fragmented Australian forests. In Landscape Ecol 29 (4), pp. 745–758. DOI: 10.1007/s10980-014-0015-1.

Villéger, Sébastien; Mason, Norman W. H.; Mouillot, David (2008): New multidimensional functional diversity indices for a multifaceted framework in functional ecology. In Ecology 89 (8), pp. 2290–2301. DOI: 10.1890/07-1206.1.

Wang, Gang; Harpole, Clifford E.; Trivedi, Amit K.; Cassone, Vincent M. (2012): Circadian regulation of bird song, call, and locomotor behavior by pineal melatonin in the zebra finch. In Journal of biological rhythms 27 (2), pp. 145–155. DOI: 10.1177/0748730411435965.

Wilsey, Brian J.; Chalcraft, David R.; Bowles, Christy M.; Willig, Michael R. (2005): RELATIONSHIPS AMONG INDICES SUGGEST THAT RICHNESS IS AN INCOMPLETE SURROGATE FOR GRASSLAND BIODIVERSITY. In Ecology 86 (5), pp. 1178–1184. DOI: 10.1890/04-0394.

Zsebők, Sándor; Schmera, Dénes; Laczi, Miklós; Nagy, Gergely; Vaskuti, Éva; Török, János; Zsolt Garamszegi, László (2021): A practical approach to measuring the acoustic diversity by community ecology methods. In Methods Ecol Evol 12 (5), pp. 874–884. DOI: 10.1111/2041-210X.13558.

