## Supplementary Material for "A framework for the quantification of soundscape diversity using Hill numbers"

### S.1: Assessing the effect of sampling duration and sampling regime on soundscape diversity metrics and their observed relationship with richness of soniferous species

In this section, we assessed the effect of sampling duration (i.e., the number of full sampling days (24h) in the recording period) and sampling regime (i.e., the temporal schedule that is used to record the soundscape throughout the recording period) on the soundscape diversity metrics described in the main text. Additionally, we were interested in how these two factors influenced the observed relationship between the soundscape richness and the richness of soniferous species. Our aim here was to offer recommendations regarding sampling design using the soundscape diversity workflow described in this work and provide a framework for rarefaction/extrapolation in case of unequal sampling size between sites.

To address these questions, we used the same set of plots as described in the empirical case study (main text – section 3), for which 4-9 full days of soundscape recordings were acquired using a 1 / 5 min sampling regime at a 44.1 kHz sampling rate in Brazilian Amazonia. To assess the effect of the sampling regime, we subsetted the obtained OSU-by-sample matrix for each plot using the following recording schedules: 1 min / 10 min; 1 min / 15 min; 1 min / 20 min; 1 min / 30 min; 1 min / 60 min. For sampling sites containing multiple plots, the obtained OSU-by-sample incidence matrices were grouped across plots.

#### *1.1. A protocol for sampling effort equalization using iNEXT*

To simulate the soundscape diversity metrics for a range of sampling durations, we can use the obtained OSU-by-sample incidence matrices to compute sample-size-based rarefaction/extrapolation curves for each site at multiple orders of diversity using the R-package iNEXT (Hsieh et al. 2016). This package provides a range of functions for the computation of Hill numbers from raw incidence data and rarefaction and extrapolation with 95% confidence intervals. If the goal of the study is to simply rank the soundscape diversity of multiple communities, the sample extrapolation size can be extended several times the observed sample size (Chao and Jost 2012). Yet, if the goal is to estimate exact relationships between communities, Chao and Jost (2012) recommend extrapolation to double the observed reference sample size at most. In the empirical case study described in the main text, we wanted to quantify the exact relationships in the soundscape diversity metrics for a set of sites along a gradient in the richness of soniferous species (birds, anurans, and primates). As the minimal sample size is four full days, we interpolated/extrapolated the soundscape diversity metrics to eight days of sampling (twice the minimal observed reference sample size) for each site and sampling regime. This way, we accounted for unequal sampling effort among sites while retaining the maximum amount of information in the datasets.

#### *1.2. Providing recommendations regarding the sampling duration*

##### *1.2.1. The impact of sampling duration and regime on the relative soundscape richness ranking between sites*

Using our workflow, we are interested in providing accurate quantification and comparison of soundscape diversity values for sites in a landscape. As such, we hope to provide recommendations regarding what constitutes an ideal in-field sampling effort to reliably quantify relationships between sites. Yet, the sampling duration can only be reliably extrapolated to double the minimal sampling effort to quantify the exact relationships among sites. Given the constraints of our dataset, as the minimal sampling duration is 4 days, we could not reliably assess how longer sampling durations (e.g., >20 days) influence these exact relationships. Instead, we focussed on the relative soundscape richness ranking among islands, which can be extrapolated to multiple times the minimal sampling effort in a reliable manner (Chao and Jost 2012).

To do so, we investigated the change in the soundscape richness ranking among islands along a sampling effort gradient for a set of sampling regimes. Specifically, we computed the soundscape richness of each island and sampling regime for a set of sampling effort values ranging from 1-28 days. Next, we calculated the richness ranking among islands at each sample effort value (i.e. the number of 24h sampling days). To quantify at which sampling effort the richness ranking stabilises among islands, thus providing the most accurate picture of the relationship among sites in the landscape, we computed the Mean Rank Shift (MRS) value using the '*codyn'* R-package (v2.0.5; Hallett et al. 2016). This metric quantifies the average change in a ranked list between two consecutive periods. Finally, to elucidate at which sampling duration the Mean Rank Shift approaches a zero asymptote, we fitted negative exponential models (y ~ a * exp(-b * x)), using the *'SSasymp'* function for selfstarting models from the '*stats*' R-package (R Core Team 2020) to obtain starting values, and the *'nls'* package (v1.0-2; Baty et al. 2015) to fit the models (Fig. S1; Table S1).

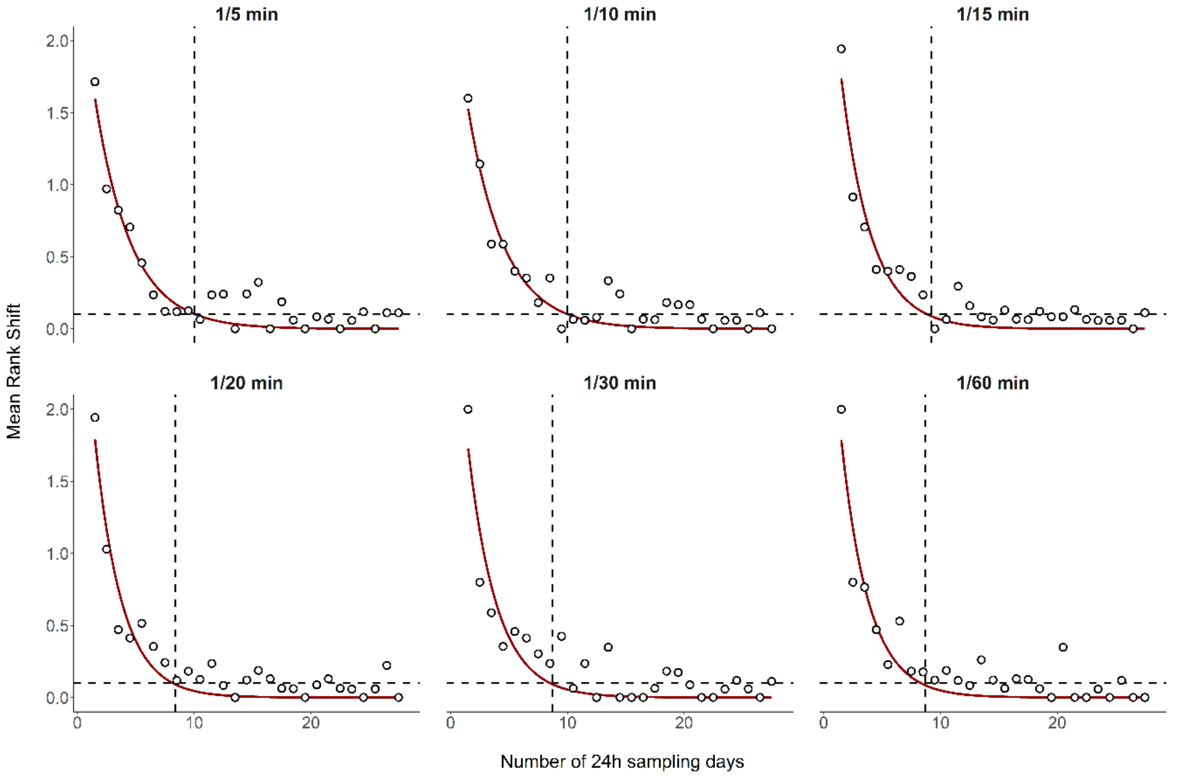


***Figure S1:*** *A set of scatterplots displaying the relationship between the sampling effort (number of 24h sampling days) and the Mean Rank Shift in the soundscape richness values for 35 sites and six sampling regimes in an Amazonian rain forest landscape. The maroon line represents a negative exponential model approaching an asymptote zero (y ~ a * exp(-b * x)). The horizontal dashed lines represent the point at which the function start approaching the zero asymptote (Mean Rank Shift = 0.1).*

***Table S1:*** *Summary of the negative exponential models (y ~ a * exp (-b * x)) fitted for different sampling regimes, in which y represents the Mean Rank Shift and x represents the number of full sampling days (p < 0.001 for all fitted models).*

| Sampling regime | parameters | value | SE |
|---|---|---|---|
| **1 / 5** | a | 2.5906 | 0.27159 |
| | b | 0.3225 | 0.03476 |
| **1 / 10** | a | 2.4595 | 0.26169 |
| | b | 0.3182 | 0.03495 |
| **1 / 15** | a | 3.1125 | 0.35674 |
| | b | 0.3877 | 0.04295 |
| **1 / 20** | a | 3.4369 | 0.42608 |
| | b | 0.4344 | 0.04985 |
| **1 / 30** | a | 3.1790 | 0.50778 |
| | b | 0.4061 | 0.06164 |
| **1 / 60** | a | 3.3756 | 0.48392 |
| | b | 0.4240 | 0.05681 |

We found that the negative exponential models fit the data well (Fig. S1). Moreover, the parameter values were all significantly different from zero and have low standard errors. Based on these models, the Mean Rank Shift among islands seemed to approach an asymptote for MRS ~ 0.1 at approximately 8-10 sampling days for all sampling regimes. As such, knowing that the sampling effort can be reliably extrapolated to double the minimum reference sample size, we recommend future studies looking for the exact relationship in the soundscape diversity between sites attempt to record the soundscape for a minimum of 5 days per site.

Given that the sampling regime employed in this study (1 min / 5 min) does not correspond with regimes used in other studies assessing the required sampling effort (*e.g.* continuous sampling), we cannot directly compare the number of required sampling days to capture the soundscape reliably. Instead, we will use the total number of sampling hours per site as an indicator of the required effort. We deem this a good proxy for sampling effort for two reasons: (i) the total number of hours that can be recorded per site is directly influenced by the storage space available on the acoustic sensor's memory card, a factor which is often limiting the sampling effort in field studies; (ii) the total number of hours can be directly compared for studies with differing sampling regimes. In our study, the soundscape richness ranking among sites stabilized after 8-10 sampling days. Knowing that our most intense sampling regime was 1 min / 5 min, this corresponds to 38.4-48 hours of recording per site. This recording duration associated with the optimal sampling duration is considerably lower than other minimal sampling durations previously reported in the literature (e.g. 120 hours in Bradfer-Lawrence et al. 2019). If the sampling design is determined by the available storage on the acoustic sensor's memory card, this is good news, as cards with a lower capacity could be used or the excess storage space could be allocated to sampling the soundscape at higher sampling rates rather than for more days or higher intensity regimes.

#### *1.2.2. The impact of sampling regime on the observed soundscape richness – taxonomic richness relationship*

In addition to the soundscape richness rank change in function of sampling effort and regime, we investigated how the sampling regime influences the soundscape richness – taxonomic richness relationship among islands for a fixed sampling effort (number of 24h recording days). As we were now interested in the exact relationship between islands, and since we wanted to account for unequal sampling effort between islands, we interpolated/extrapolated the sampling effort to 8 days (twice the minimal sampling effort) using iNEXT (Hsieh et al. 2016). Although this sampling duration was at the lower end of the minimally recommended duration (see S.1.2.1), an inspection of Fig. S1 revealed that at 8 sampling days, for each additional sampling day the Mean Rank Shift changed by less than 0.1 on average. Moreover, the Mean Rank Shift decreased towards the zero asymptote as more sampling days were added. As such, the change in the overall relationship between the soundscape richness and richness of soniferous species at longer sampling durations was likely minimal.

For each of the sites, after rarefaction/extrapolation, we computed the soundscape richness and assessed the relationship with the richness of soniferous species. As the sampling regime influences the number of OSUs which can be detected in functional trait space, we divided the soundscape richness values by the total number of detectable OSUs in this space to get the percentage of space occupied. Finally, we computed the Pearson correlation coefficient and $R^2$-value for a simple linear regression model between the soundscape richness and the richness of soniferous species (Fig. S2).

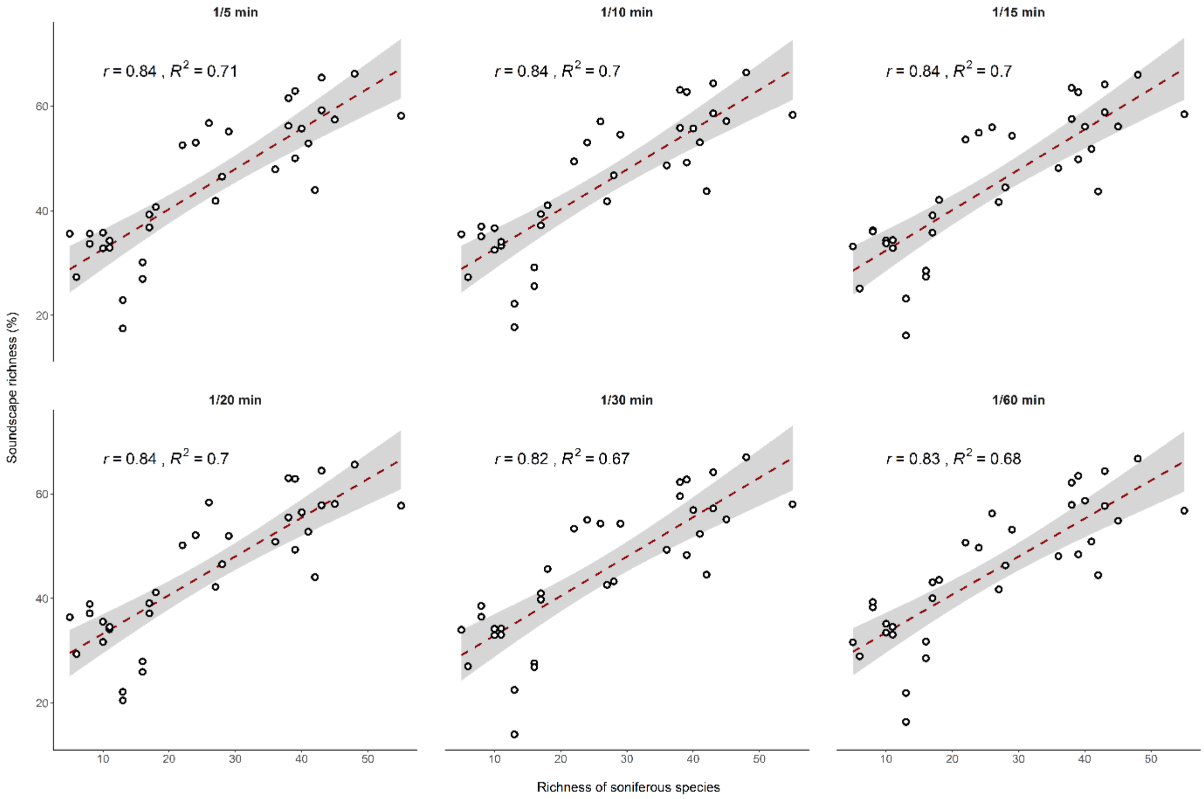


***Figure S2:*** *Relationships between the soundscape richness (as the percentage of functional trait space occupied by OSUs) and the richness of soniferous species for a wide range of sampling regimes. Pearson correlation coefficients and associated p-values, which are given for each sampling regime ($p < 0.001$ in all instances), show that the predictive power of soundscape richness is largely invariant in relation to these sampling regimes.*

The scatterplots and associated Pearson correlation coefficients (Fig. S2) revealed that, at equal sampling effort, the relationship between the soundscape richness and the richness of soniferous species remained high ($r > 0.8$) even for the sparse sampling regimes. This was contrary to previous findings reported in the literature (e.g. Bradfer-Lawrence et al. 2019), which found that intense sampling regimes (*e.g.* continuous sampling) were required to capture soundscapes reliably – suggesting the workflow described here is robust and can pick up on ecological patterns, even at low sampling regimes of lower intensity.

**S.2: Assessing the effect of window length on soundscape diversity metrics**

As with time-frequency bins in spectrograms, the resolution of OSUs in the frequency domain of functional trait space is dictated by the sampling rate and window length with which the Fast Fourier Transformation is performed. The data of the empirical case study was acquired using a 44,100 Hz sampling rate and then truncated at 14,125 Hz. As such, it is the choice of window length that dictates the resolution in the frequency domain. The choice of window length determines the sensitivity of the CVR-index values to different types of sound and depends on the soniferous community in the study area of interest. A window length of 512 (Gasc et al. 2013; Rodriguez et al. 2014; Eldridge et al. 2018; Burivalova et al. 2018; Phillips et al. 2018), 256 (e.g. Campos-Cerqueira et al. 2020) and 1024 samples (e.g. Machado et al. 2017) have all been used to capture the audible soundscape in a range of environments.

Here, we investigated the effect of window length choice on the proposed soundscape diversity metrics and their relationship with the richness of soniferous species in the landscape. To do so, we computed the CVR-index for each one sound minute file using a sampling rate of 44,100 Hz and a window length of 128, 256, 512 and 1024 samples. For each of these window lengths, we applied the same analytical workflow as described in the main body of this manuscript. We calculated the soundscape richness and assessed its relationship with the richness of soniferous species using the Pearson correlation test and $R^2$-value for a simple linear regression model.

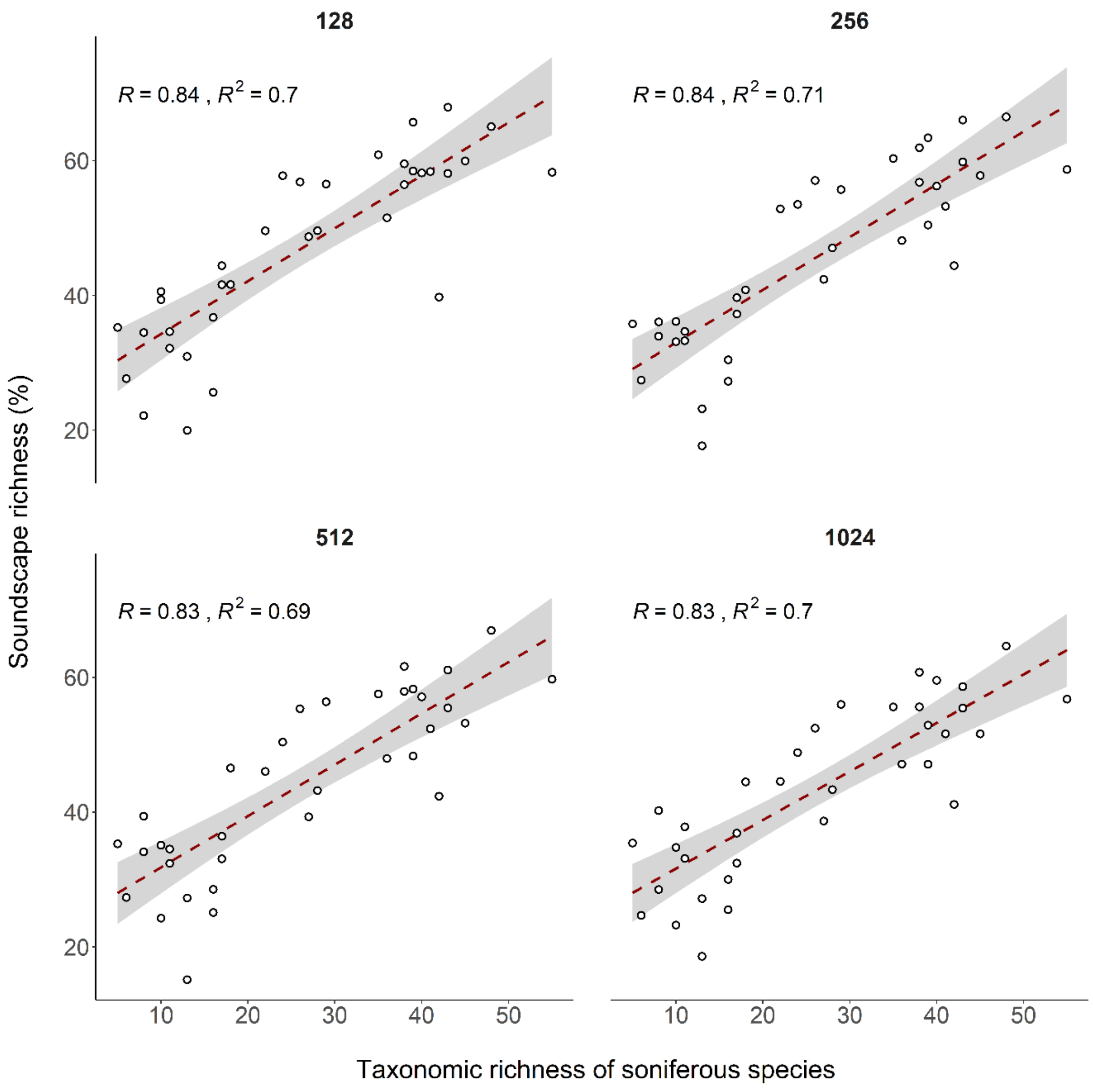


***Figure S3:*** *A set of scatterplots showing the relationship between the soundscape richness (as the percentage of the total number of detectable OSUs) and the richness of soniferous species for a set of window lengths (wl = 128; 256; 512; 1024). For each window length, the Pearson correlation coefficient (r) and* $R^2$*-value are provided (*$p < 0.001$ *in all instances).*

We found that the choice of window length had a negligible impact on the observed relationship between the soundscape richness and the richness of soniferous species (Fig. S3) – suggesting the proposed soundscape richness metric is sensitive to ecological patterns regardless of these methodological variations. For future studies, we advise the use of a 256-sample window length, as this has been previously used in the literature and provides a good correlation with the richness of sound-producing organisms.

### S.3: Assessing the effect of threshold choice on the soundscape richness – taxonomic richness relationship

Choosing the binarisation threshold which is used to obtain detection / non-detection values for each OSU in the 24-hour functional trait space constitutes an important step in our workflow. The choice of this threshold value depends on the sound transmission characteristics of the habitat under investigation, and the amount and type of background noise in the environment. The ideal threshold value removes the influence of low-amplitude and transient or short-duration noise on the soundscape diversity metrics described in this study, thus increasing the sensitivity to the biophonic taxonomic richness. Several thresholding approaches exist to achieve this objective. For instance, in Burivalova et al. (2018), a range of amplitude threshold values were trialled, looking for the value which yielded a near-normal distribution of the variable of interest across all sites in the study. In Aide et al. (2017), a fixed threshold value was used across all sites to determine the presence of sound. Here, we investigated various approaches and how they influenced the observed relationship between the soundscape richness and the richness of soniferous species. The preferred thresholding method is the one that increases the sensitivity of the proposed metrics to the richness of soniferous species.

#### *3.1. Applying a constant threshold value across all sites*

For our first approach, we applied a constant threshold value for all sites in the study. We applied the same analytical workflow as described in S.1, however, to assess which constant threshold value yields the best relationship between the soundscape richness and the richness of soniferous species, we trialled a range of constant threshold values between 0.01 and 0.5 at 0.1 intervals. For each of these threshold values, we calculated the soundscape richness as described in the main text. Finally, per Burivalova et al. (2018), we assessed which of the threshold values yielded the best near-normal distribution of soundscape richness values, and which of the threshold values yielded the best correlation with the richness of soniferous species.

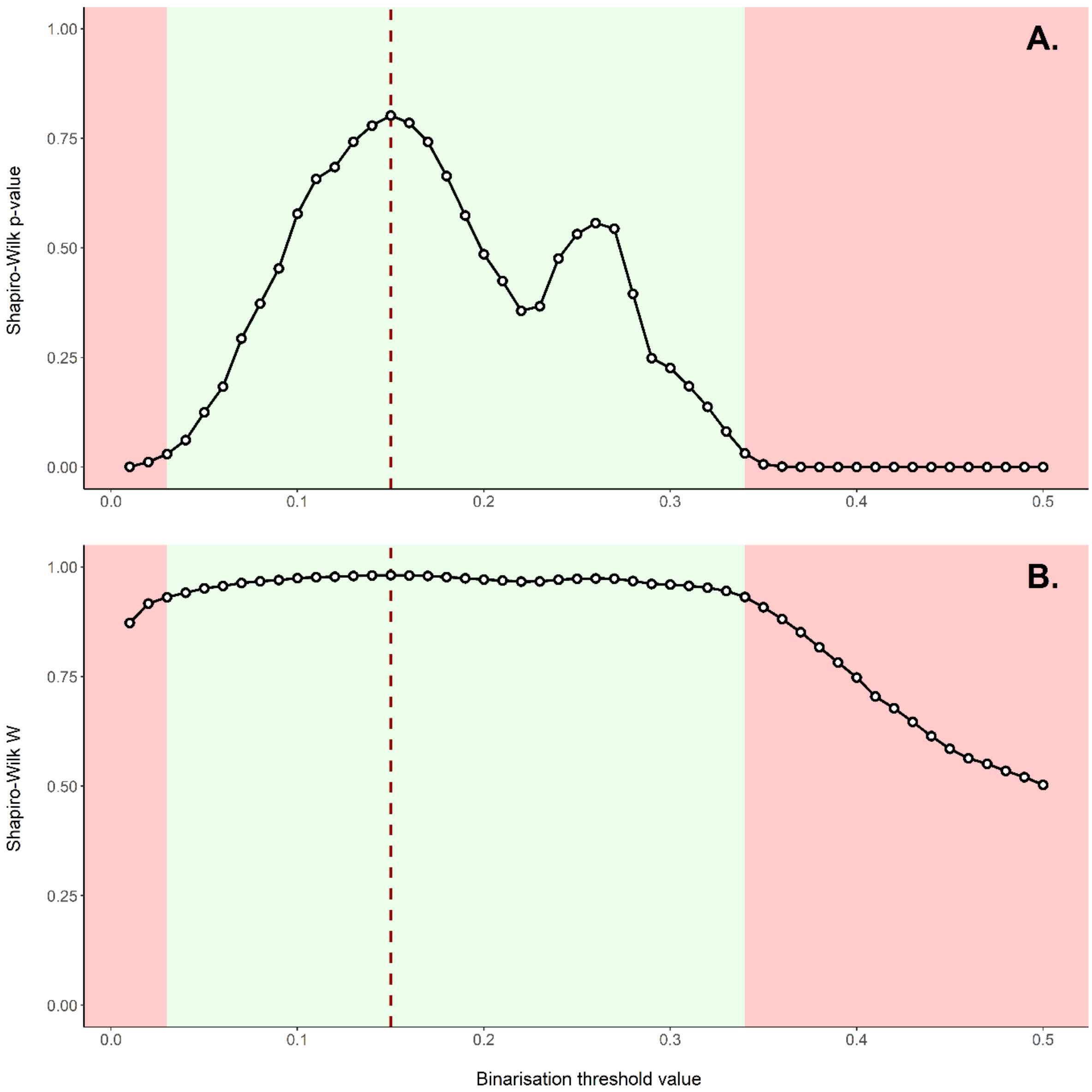


***Figure S4: A.*** *The relationship between the binarisation threshold value and the p-values acquired from the Shapiro-Wilk normality test performed on the soundscape richness data obtained at each threshold value;* ***B.*** *The relationship between the binarisation threshold value and the W-values acquired from the Shapiro-Wilk normality test performed on the soundscape richness data obtained at each threshold value. The dashed red line represents the threshold value for which the resulting soundscape richness values are most normally distributed (highest p- and W-values). The shaded red area represents the threshold values for which the resulting soundscape richness values are not normally distributed (p-value < 0.05), whereas the shaded green area represents threshold values resulting in normally distributed soundscape richness values.*

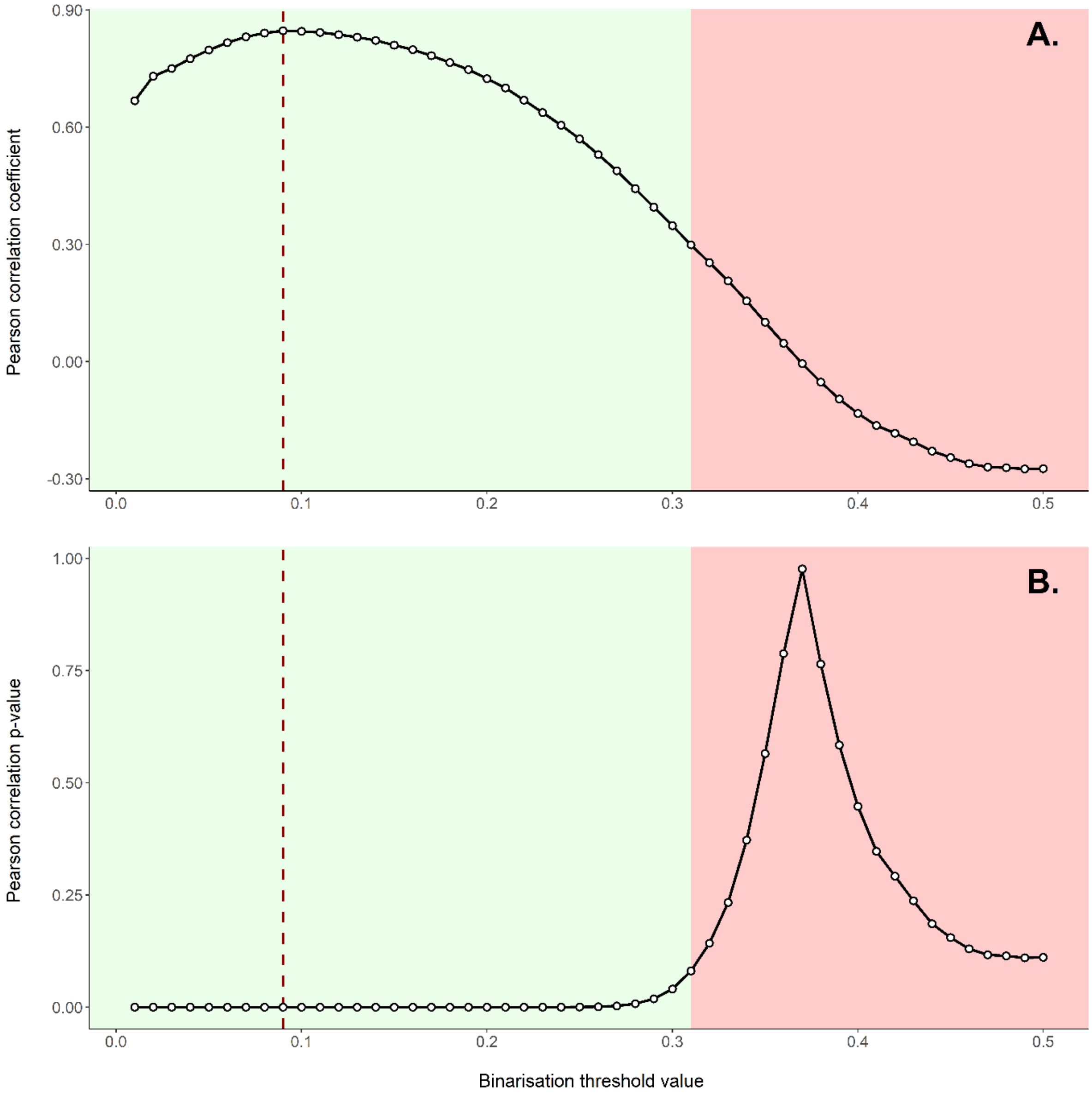


***Figure S5: A.*** *The relationship between the binarisation threshold value and Pearson's correlation coefficient (r) obtained from the Pearson correlation test between the resulting soundscape richness and the richness of soniferous species.;* ***B.*** *The relationship between the binarisation threshold value and the p-value obtained from the Pearson correlation test between the resulting soundscape richness and the richness of soniferous species. The dashed red line represents the threshold value (threshold = 0.09) for which the correlation between the soundscape richness and richness of soniferous species is highest (r = 0.84). The shaded red area represents the values for which the correlation is not significant, whereas the shaded green area represents the values for which correlations are significant.*

The most normal distribution of soundscape richness values was obtained at a constant binarisation threshold of 0.18 (Figs. S4A and S4B; r = 0.77; $R^2$ = 0.59; p < 0.001), yet this value does not correspond to the binarisation threshold which yields the highest correlation with the richness of soniferous species (threshold = 0.09; *r* = 0.84; $R^2$ = 0.71; p < 0.001; Figs. S5A and S5B). In the next section, we investigate the use of site-dependent thresholding using binarisation algorithms.

#### *3.2. Applying a site-dependent threshold using binarisation algorithms*

For our second approach, instead of applying a constant binarisation threshold, we applied a site-dependent threshold to each of the plots described in S.1. To determine the threshold value for each plot, we made use of the binarisation algorithms available in the *autothresholdr'* R-package (v1.3.11; Landini et al. 2017 - for algorithm descriptions, consult: https://imagej.net/plugins/auto-threshold). Every binarisation algorithm provides a unique binarisation threshold per plot, and unlike the previous method, threshold values can be variable between plots. We omitted the following binarisation algorithms from the analysis, as they were not suitable for our type of data: “Intermodes”, “MaxEntropy”, “Minimum”, “Yen”. Finally, we assessed which of the binarisation algorithms produced the best relationship between the soundscape richness and the richness of soniferous species using the Pearson correlation coefficient and $R^2$-value for a simple linear regression model.

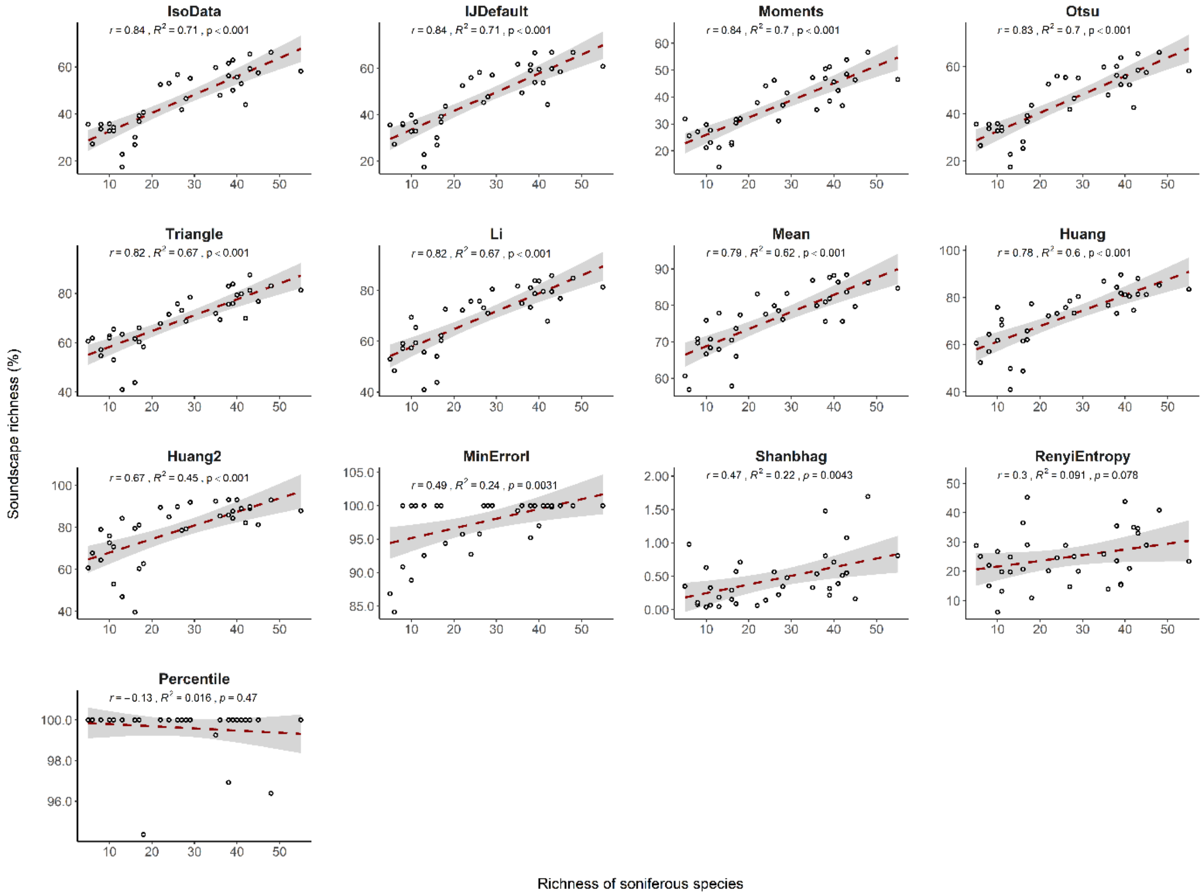


***Figure S6:*** *A set of scatterplots displaying the relationship between the soundscape richness (as the percentage of the total number of detectable OSUs) and the richness of soniferous species for the various binarisation algorithms available in the ‘autothresholdr’ R-package (Landini et al. 2017). The Pearson correlation coefficient (r) and $R^2$-values are given for each binarisation algorithm.*

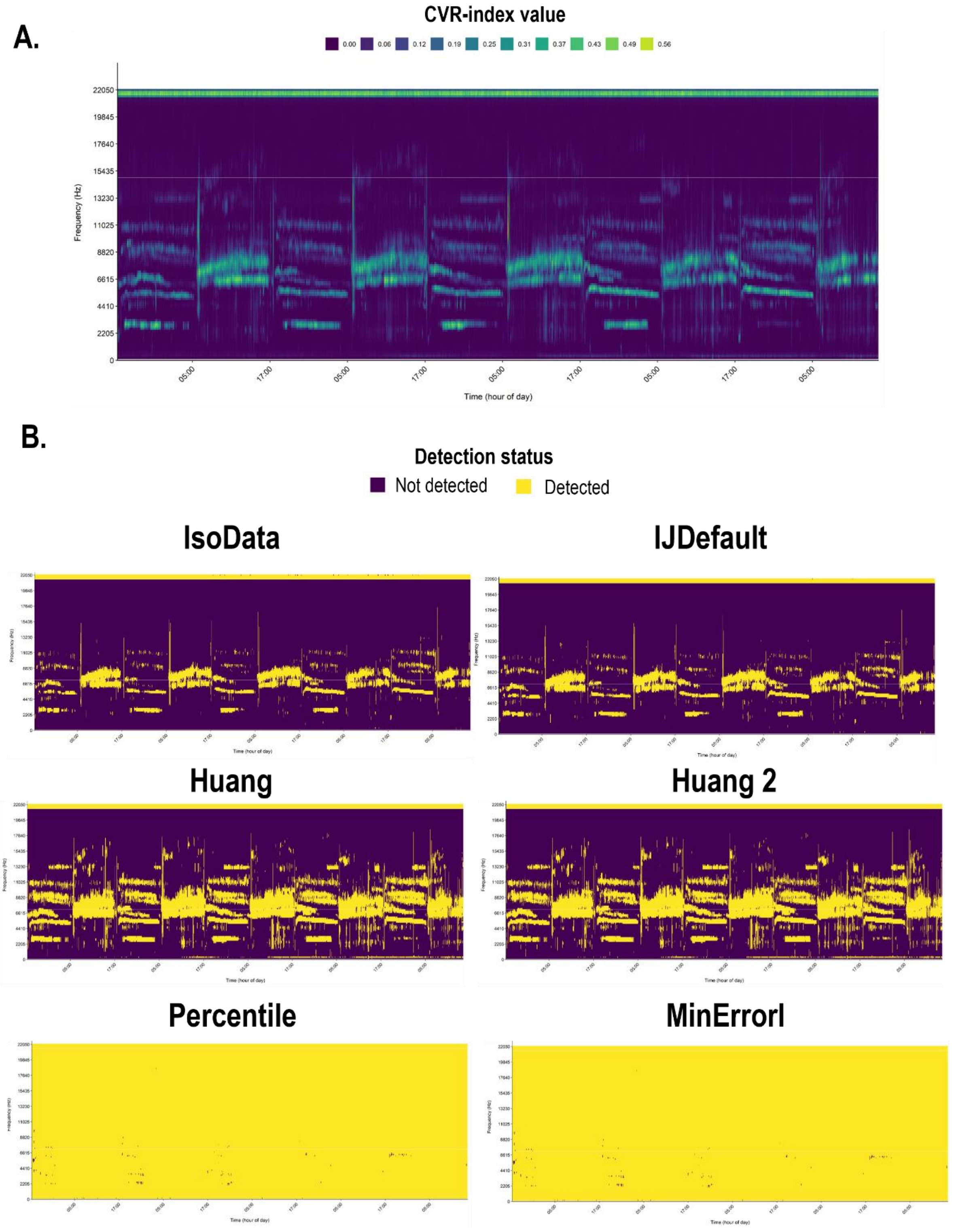


***Figure S7:*** *A visual representation of the pre- (**A.**) and post-binarisation (**B.**) soundscapes using a subset of binarisation methods available in the 'autothresholdr' R-package (Landini et al. 2017) for one of the sites in the study period. Post-binarisation plots are ranked from high ("IJDefault" and "IsoData"), to medium ("Huang" and "Huang 2") and low ("Percentile" and "MinErrorI") correlation with taxonomic richness. Visual inspection of pre- and post-binarisation plots provides insight into how the acoustic structure is captured and which threshold stringency results in good correlation with taxonomic richness.*

We found high Pearson correlation coefficients ($r > 0.80$) for the relationship between the soundscape diversity and the richness of soniferous species using the 'IsoData', 'IJDefault', 'Moments', 'Otsu', 'Triangle' and 'Li' binarisation algorithms (Fig. S6), the value of which as similar to the correlation coefficients found for constant binarisation thresholds (Fig. S5). A visual inspection of the pre- and post-binarisation plots (Fig. S7) revealed that the more stringent thresholding methods (*e.g "IsoData" and "IJDefault")*, where more sound is removed, resulted in higher correlations with taxonomic richness.

The binarisation algorithms produce a unique binarisation threshold per plot that is determined by the distribution of CVR-values in the acoustic trait space, which in turn is influenced by the soniferous community and sound transmission characteristics of the habitat. As such, these binarisation algorithms generate context-aware threshold values. Conversely, although we found that a thresholding value of around 0.1 worked well for the plots in our study, other habitats with differing sound transmission characteristics, noise levels, or acoustic communities might have a different optimal threshold value. As such, since the threshold value for binarisation algorithms is determined directly by the acoustic fingerprint of the plot, we recommend using one of the binarisation algorithms highlighted above. Further research in a wider variety of habitats is needed to confirm that the 'IsoData' algorithm performs the best consistently.

**S.4. A framework for decomposing soundscape diversity into its alpha, beta, and gamma components**

In addition to quantifying the soundscape richness (${}^{0}D$), diversity (${}^{q}D$ with $q = 1, 2, ...$), and evenness (${}^{2}D\,/\,{}^{0}D$) components, the workflow proposed in this manuscript can be used to decompose the regional metacommunity diversity (γ-diversity) into its local diversities (α-diversity) and a community turnover component (β-diversity). Moreover, it can be used to generate several measures of similarity or dissimilarity and overlap. Here, we outlined the theoretical framework for decomposing the soundscape diversity into its alpha, beta, and gamma components. In S.5.5, we provided a working example of this theoretical framework in practice.

The framework of Hill numbers follows a multiplicative relationship to decompose the soundscape diversity into its various components (Eqn. 1):

$$(1)\quad {}^{q}D_{\gamma} = {}^{q}D_{\alpha} \times {}^{q}D_{\beta}$$

$$(2)\quad {}^{q}D_{\alpha} = \frac{1}{N}\left\{\sum_{i=1}^{S}\sum_{j=1}^{N}(w_j p_{ij})^{q}\right\}^{\frac{1}{(1-q)}}$$

$$(3)\quad {}^{q}D_{\gamma} = \left\{\sum_{i=1}^{S}\left(\sum_{j=1}^{N}(w_j p_{ij})\right)^{q}\right\}^{\frac{1}{(1-q)}}$$

$$(4)\quad {}^{q}D_{\beta} = \frac{{}^{q}D_{\gamma}}{{}^{q}D_{\alpha}}$$

Here, $N$ refers to the total number of sub-systems (soundscapes), $j$ refers to each individual sub-system, and $w_j$ represents the relative weight given to each sub-system in the system. If all soundscapes are weighted equally, $w_j$ equals $1/N$. The alpha diversity is the Hill number of the averaged basic sums of the soundscapes (Eqn. 2). The gamma diversity is computed by taking the average of the relative abundance of each OSU across the soundscapes in the system and calculating the Hill number of the pooled system (Eqn. 3). The beta diversity captures the degree of heterogeneity in the OSU composition across sites (Eqn. 4). It ranges from 1 to $N$ and quantifies the relationship between the regional and local diversity, that is, how many times more diverse is the whole system in the effective number of OSUs compared to the sub-systems on average (Alberdi and Gilbert

2019). The beta diversity can also be seen as the effective number of completely distinct soundscapes in the system (Tuomisto 2010).

The framework of Hill numbers also allows us to define several measures of similarity between soundscapes in the wider system. Because beta diversity ranges between 1-*N*, it is not independent of the number of soundscapes in the system, and can thus not be used as a measure of similarity directly (Alberdi and Gilbert 2019). Instead, to compare the relative compositional difference between soundscapes across multiple systems with a different number of soundscapes, some simple transformations can be performed on the beta diversity to remove the dependence on the number of soundscapes (Jost 2007; Chao et al. 2012; Chiu et al. 2014; Alberdi and Gilbert 2019).

$$(5)\quad C_{qN} = \frac{\left[\left(\frac{1}{{}^{q}D_{\beta}}\right)^{(q-1)} - \left(\frac{1}{N}\right)^{(q-1)}\right]}{\left[1 - \left(\frac{1}{N}\right)^{(q-1)}\right]}$$

$$(6)\quad U_{qN} = \frac{\left[\left(\frac{1}{{}^{q}D_{\beta}}\right)^{(1-q)} - \left(\frac{1}{N}\right)^{(1-q)}\right]}{\left[1 - \left(\frac{1}{N}\right)^{(1-q)}\right]}$$

Equations (5) and (6) are measures of overlap between soundscapes. The local or Sørensen-type overlap (CqN) quantifies the effective average proportion of a soundscape's OSUs which are shared across all soundscapes (Chiu et al. 2014). It captures the overlap between soundscapes from the sub-system's perspective (Alberdi and Gilbert 2019). For *N* soundscapes each having *S* equally common OSUs and sharing *A* OSUs between them, this function reduces to $C_{qN}=A/S$. The regional or Jaccard-type overlap ($U_{qN}$) quantifies the effective proportion of shared OSUs in a pooled assemblage of soundscapes, and thus captures the overlap between soundscapes from a regional perspective. Assume *N* soundscapes in a region with *S* unique and equally abundant OSUs. Here, *R* OSUs are shared between all soundscapes and the remaining OSUs (*S-R*) are distributed evenly among *N* soundscapes. In this scenario, Eqn. 8 reduces to $U_{qN} = R/S$.

$$(7)\quad V_{qN} = \frac{\left(N - {}^{q}D_{\beta}\right)}{(N-1)}$$

$$(8)\quad S_{qN} = \frac{\left(\frac{1}{{}^{q}D_{\beta}} - \frac{1}{N}\right)}{\left(1 - \frac{1}{N}\right)}$$

Equations (7) and (8) are measures of turnover in OSUs between soundscapes (Harrison et al. 1992; Jost 2007). The local or Sørensen-type turnover complement ($V_{qN}$) quantifies the normalised OSU turnover rate with respect to the average soundscape (Alberdi and Gilbert 2019). It measures the proportion of a typical soundscape which changes as one goes from one soundscape to the next (Harrison et al. 1992; Chao et al. 2012; Jost 2007). The regional or Jaccard-type turnover complement ($S_{qN}$) quantifies the proportion of the regional soundscape diversity contained in the average assemblage and is a measure of regional homogeneity. All of the aforementioned similarity indices can be transformed into metrics of dissimilarity by taking their one-complement ($1 - X_{qN}$) (Alberdi and Gilbert 2019).

### S.5: Simulating artificial soundscapes to demonstrate the soundscape diversity workflow's desirable properties

In this section, we used simulated artificial soundscapes to construct simple examples which illustrate that the proposed soundscape diversity framework abides by a set of fundamental criteria for functional diversity indices and behave in an ecologically intuitive manner. To do so, we adopted some of the criteria relevant to our workflow outlined in Ricotta (2005), Villéger et al. (2008) and Mouchet et al. (2010), and added to these some key behaviours we deemed fundamental for our workflow to work as required (Table S2). As suggested in Villéger et al. (2008), here it is not important that each index matches each criterion, but rather that the ensemble of the indices does. In section S.5.1, we discussed the criteria which can be confirmed without a need for explicit testing. Further on, in sections S.5.2 – S.5.6, we used simulated artificial soundscapes to prove that the workflow abided by the fundamental properties we desire.

***Table S2:*** *A summary of some desired criteria and properties for the soundscape diversity indices described in this workflow. A green tick mark indicates the index follows the criterion, whereas a red tick mark indicates it does not. For criterion 4, two symbols are provided, indicating whether that soundscape diversity metric follows this criterion for the closest other metric in the table, looking from left to right. For instance, the soundscape richness is not independent from the soundscape diversity at q=2 but is independent from the soundscape evenness.*

| **Criterion** | **Soundscape richness ($^0D$)** | **Soundscape diversity ($^1D$, $^2D$, …)** | **Soundscape evenness ($^2D$ / $^0D$)** |
|---|---|---|---|
| (1) The indices can only have positive values (see section 5.1) | ✓ | ✓ | ✓ |
| (2) The indices can be strictly contained between 0 – 1 (see section 5.1) | ✓ | ✓ | ✓ |
| (3) The indices are independent of the species richness (see section 5.1) | ✓ | ✓ | ✓ |
| (4) The indices are independent of one another (see section 5.3) | ✗ ✓ | ✗ ✗ | ✓ ✗ |
| (5) The indices are monotonous (see section 5.2) | ✓ | ✓ | ✗ |
| (6) The indices abide by the replication principle (see section 5.4) | ✓ | ✓ | ✗ |
| (7) The indices can be decomposed into alpha, beta, and gamma components (see section 5.6) | ✓ | ✓ | ✗ |

#### *5.1. Fundamental properties of functional diversity indices*

Several of the desired properties of our soundscape diversity workflow were fulfilled because of how the workflow and diversity indices have been set up, and thus did not need explicit testing. For instance, all our indices were strictly positive and could be constrained between 0-1. The soundscape richness and diversity (at $q$ = 1, 2, ...) could be constrained between 0-1 by dividing by the diversity value by the maximum possible detectable OSUs in the acoustic functional trait space. The soundscape evenness values were constrained between 0-1 by default. Moreover, the criterion of independence from the species richness was true for all indices, as we did not use any taxonomic information in our workflow. As such, any potential correlation between the diversity measures and the taxonomic richness stemmed from underlying processes of species assembly, and not inherent correlations stemming from how the metrics are computed. In the next section, we tested the remaining criteria and behaviours using simulated soundscapes.

#### *5.2. The soundscape richness, evenness and diversity are monotonous*

The criterion of monotonicity states that a subset of the community should always have a lower diversity value than the total community. To test this, we created 100 simplified simulated soundscapes sampled using a 1 min / 5 min sampling regime (resulting in 288 temporal bins in a 24h period), and a 0 – 11,025 Hz frequency domain generated using a 256-frame window length (resulting in 64 frequency bins). In this case, the total number of detectable OSUs was 18,432 (288 * 64). We considered each soundscape to be completely filled, having a soundscape richness of 18,432 OSUs. For every simulated soundscape, the relative abundance values of OSUs were sampled so that the number of highly abundant OSUs in the community (relative abundance = 1) ranges from 1-100% of OSUs – the remainder of OSUs being rare (relative abundance = 0.01). By doing so, we altered the proportion of dominant species in the community, thus creating one hundred evenness classes for each soundscape. For each of these 100 soundscapes, we subsetted the soundscape using sample sizes ranging between 1-99% of the total soundscape richness at 1% increments. Then, for each sample size, we randomly sampled the correct number of OSUs 100 times, resulting in a total of 1,000,000 replicates (100 soundscapes with different evenness values * 100 subset sizes * 100 replicates). We calculated the soundscape richness, diversity ($q$=2), and evenness ($^2D / {^0D}$) for each soundscape. To test whether each of the metrics abides by the criterion of monotonicity, we subtracted the richness, diversity, and evenness metrics for each subsetted soundscape from their respective original soundscape metrics. If any negative values are present in the data, this suggests the soundscape diversity metric of the subsetted soundscape was larger than that of the original soundscape, and thus the criterion of monotonicity does not hold.

As expected, the monotonicity criterion held true for the soundscape richness and soundscape diversity (Fig. S8), as a subset of OSUs could never be richer than the total pool. For the evenness, however, the criterion of monotonicity did not hold, as a subset of OSUs could result in a higher evenness value than the total pool if the relative abundances of the subset were more even (Fig. S8).

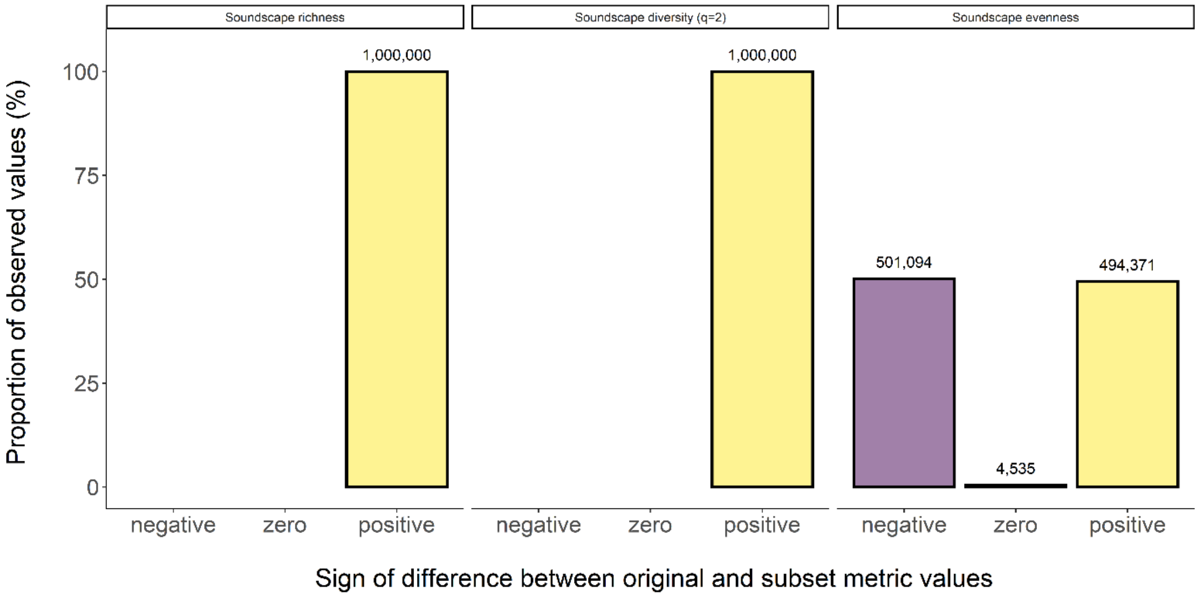


***Figure S8:*** *A barplot displaying the proportion of observations that had a negative, zero or positive sign for the difference between the soundscape richness, diversity (q=2) and evenness of the original soundscape and the respective metric of the subsetted soundscape. Negative values indicate the value of the subset metric was larger than the original value, and thus the metric does not abide by the criterion of monotonicity. Above each bar, the raw number of observations for that sign is displayed.*

#### *5.3. Soundscape diversity metrics are independent of one another*

For our next criterion, we wanted to assess whether our soundscape diversity metrics are strictly independent of one another, capturing unique aspects of acoustic trait space usage. To test this, we generated simulated soundscapes with randomised variation in the functional trait values. We considered the same simplified functional trait space as in section 5.2, consisting of 18,432 detectable OSUs. We considered one hundred soundscape richness classes (1-100% of OSUs detected). For each of these richness classes, the equivalent number of OSUs in functional trait space was generated. For instance, for the functional trait space with 18,432 potential OSUs, the 10% richness class resulted in 0.1*18,432 = 1843 OSUs. As before, for each richness class, the relative abundance values of OSUs were sampled so that the number of highly abundant species in the community (relative abundance = 1) ranges from 1-100% of OSUs – the remainder of OSUs being rare (relative abundance = 0.01). As such, we have created 10,000 replicates along the richness-evenness range. The soundscape richness, diversity and evenness were computed for these 10,000 datasets, and the Pearson correlation coefficient and $R^2$-value for a simple linear regression model were calculated for each combination of indices to assess independence. We expected the soundscape richness and evenness to be strictly independent of one another, capturing unique aspects of the soundscape. Conversely, as the soundscape diversity at $q$=2 incorporates both aspects of the soundscape richness and evenness, we expected it to be positively correlated with an increasing soundscape richness and evenness.

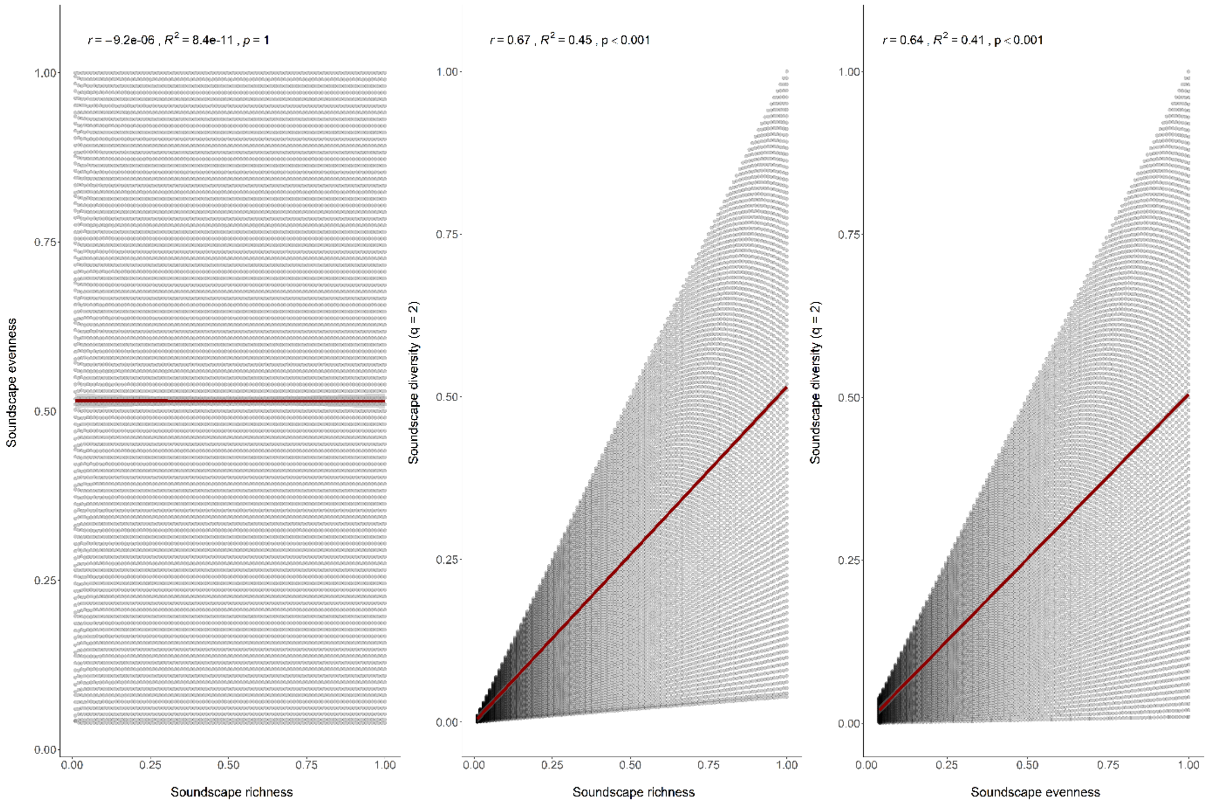


***Figure S9:*** *The relationship between the soundscape diversity variables (soundscape richness, evenness, and diversity) for 10,000 simulated soundscapes with richness values ranging between 1-100% of the total number of detectable OSUs, and relative abundance values sampled so that the evenness value covers the range from 0-1. The Pearson correlation coefficient ($r$ < 0.001) and associated $R^2$-value ($R^2$ < 0.001) reveal there is no relationship between the soundscape richness and evenness, confirming they are strictly independent. The soundscape diversity at q = 2 is positively correlated with both the soundscape richness ($r$ = 0.67) and the soundscape evenness ($r$ = 0.64).*

The Pearson correlation coefficient and associated $R^2$-value obtained for the simulated soundscapes demonstrated that the soundscape richness and evenness were independent of one another (Fig. S9), thus satisfying our independence criterion. Moreover, as expected, the soundscape diversity at $q$=2 demonstrated a positive relationship with both the soundscape richness and evenness, being sensitive to changes in both.

#### 5.4. Soundscape metrics abide by the replication principle

For our next test, we wanted to test whether the diversity indices described in this study abide by some fundamental behaviours required for biodiversity indices. The replication principle states that for two equally diverse communities with identical relative abundance distributions and no shared species, the diversity of the pooled community assemblage should be twice as high (Hill 1973; Jost 2006). Inversely, when the number of OSUs in the system is reduced by half, so should the diversity. Yet, regular diversity indices such as the Shannon and Simpson indices do not follow this intuitive notion of diversity. For these indices, the change in index values is not proportional to the change in the underlying diversity of the system. As such, treating these diversity indices as diversity values can lead to gross misinterpretation of results (Alberdi and Gilbert 2019). Hill numbers follow the replication principle, which makes changes in their magnitude easily interpretable and ensures that the beta diversity, computed as the ratio between alpha and gamma, accurately reflects the compositional similarity of soundscapes.

Here, we demonstrated that the soundscape diversity indices (${}^{q}D$ with $q$ = 0, 1, 2, …) proposed in this study abided by the replication principle. To do so, we modified a real-life soundscape which was generated for one of the sites in section S.1, to create three artificial soundscapes. The first two artificial soundscapes we made to be equally diverse with an identical relative abundance distribution, but no shared species. To achieve this, for the OSUs occurring between 12:00h – 23:59h in the real-life soundscape, we set the incidence values to zero, thus generating a half-filled soundscape that represents our first artificial soundscape (Fig. S10 – soundscape 1). Next, to generate our second artificial soundscape with an equal diversity and relative abundance distribution but no shared OSUs, we copied the OSUs and their relative abundance values occurring between 00:00h – 11:59h to the period from 12:00h – 23:59h. Next, we set the incidence values for the OSUs occurring between 00:00h – 11:59h to zero (Fig. S10 – soundscape 2). This way, our second artificially generated soundscape contained the exact diversity and relative abundance distribution of OSUs, but no shared OSUs. For our final artificial soundscape, we pooled artificial soundscapes 1 and 2 (Fig. S10 – pooled soundscape).

To demonstrate that the diversity of the pooled assemblage is twice the diversity of the sub-soundscapes (soundscapes 1 and 2), we computed the soundscape diversity ($q$ = 0, 1, 2), as well as equivalent traditional diversity indices, the Shannon and Simpson diversity indices, for all artificially generated soundscapes.

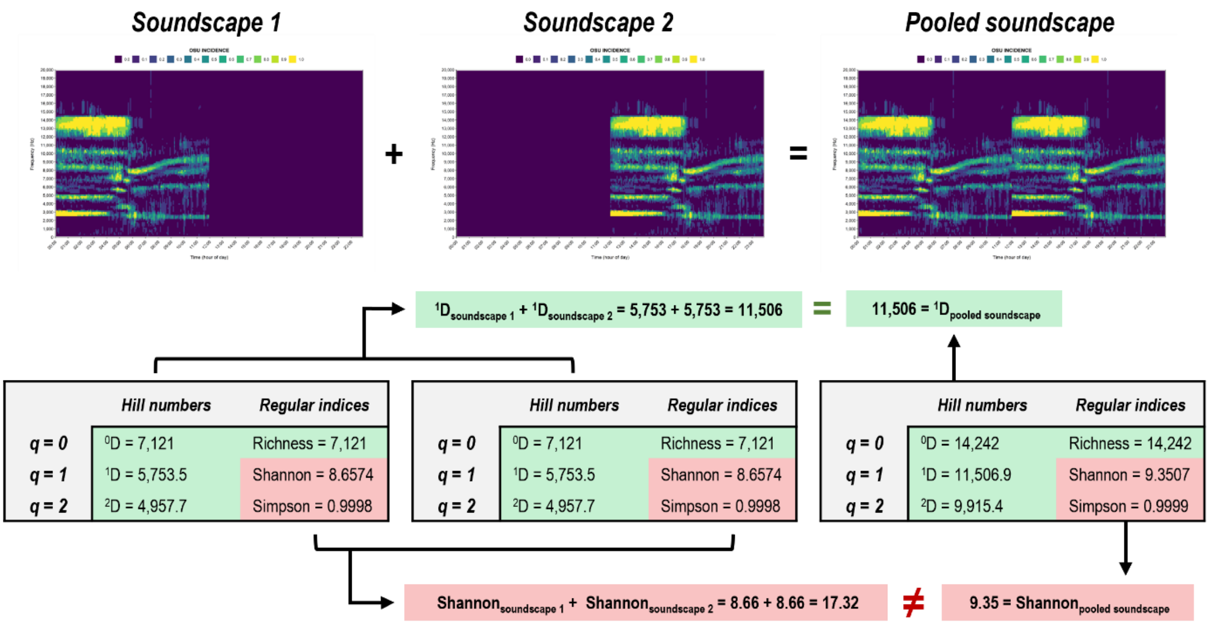


***Figure S10:*** *A visual representation of our workflow's adherence to the replication principle, demonstrated using artificial soundscapes. The replication principle states that, for two soundscapes that are equally diverse and have an identical relative abundance distribution, but without any species in common (soundscape 1 and soundscape 2), the diversity of the pooled soundscape should be twice as high. We demonstrate this holds true for the Hill numbers applied to our workflow (green shaded areas), but not for the equivalent traditional diversity indices (red shaded areas).*

Using these artificially generated soundscapes, we demonstrated that the soundscape diversity metrics proposed in this study abided by the replication principle, whereas the Shannon and Simpson indices which are commonly used in soundscape research did not.

#### 5.5. Diversity profiles

Even when diversity metrics abide by the replication principle, a single diversity metric only portrays part of the information. The perceived diversity of a soundscape depends on the importance the researcher gives to the commonness or rarity of OSUs, which is modulated by parameter *q*. For instance, one soundscape might have a higher richness but lower evenness than another – information that is lost when only looking at a single index. Diversity profiles plot Hill numbers in function of the parameter *q*, thus providing an accurate graphical representation of the shape of the acoustic community – or a type of soundscape fingerprint (Fig. S11). They provide insight into the change in perceived soundscape diversity as the emphasis shifts from rare to common OSUs, graphically illustrating the evenness and degree of dominance in the community (Leinster and Cobbold 2012; Chao et al. 2014). The left-hand side of the diversity profile yields information about the soundscape richness, valuing rare and common species equally. The right-hand side gives information about the diversity of common or dominant OSUs. Thus, they allow researchers to investigate the diversity of the soniferous community from multiple perspectives, providing a more comprehensive method for inter-soundscape comparison than any single index (Chao and Jost 2015).

We demonstrated the utility of diversity profiles by generating a set of four simulated soundscapes with an equal soundscape richness, but varying soundscape diversity and evenness. To do so, we used one of the soundscapes we previously produced in the case study and modified the relative abundance distribution of the OSUs in the community to simulate different degrees of soundscape evenness. We simulated four soundscapes along the evenness gradient by modifying the OSU abundances so that the number of highly abundant species in the community (relative abundance = 1) ranged from 25%, 50%, 75% and 100% of OSUs. The remainder of rare OSUs were obtained by randomly sampling their relative abundance along a range of 0.01-0.5 using a 0.01 interval. For each of these simulated soundscapes along the evenness gradient, we computed the soundscape diversity along a range of diversity orders $q$ from 0.01-5.0 at 0.01 intervals. Finally, to produce the soundscape diversity profiles, we plotted the soundscape diversity in function of diversity order $q$ for the four simulated soundscapes with differing evenness (Fig. S11). In addition to the soundscape diversity profile, we provided a graphical representation of the acoustic trait space for each of the simulated soundscapes.

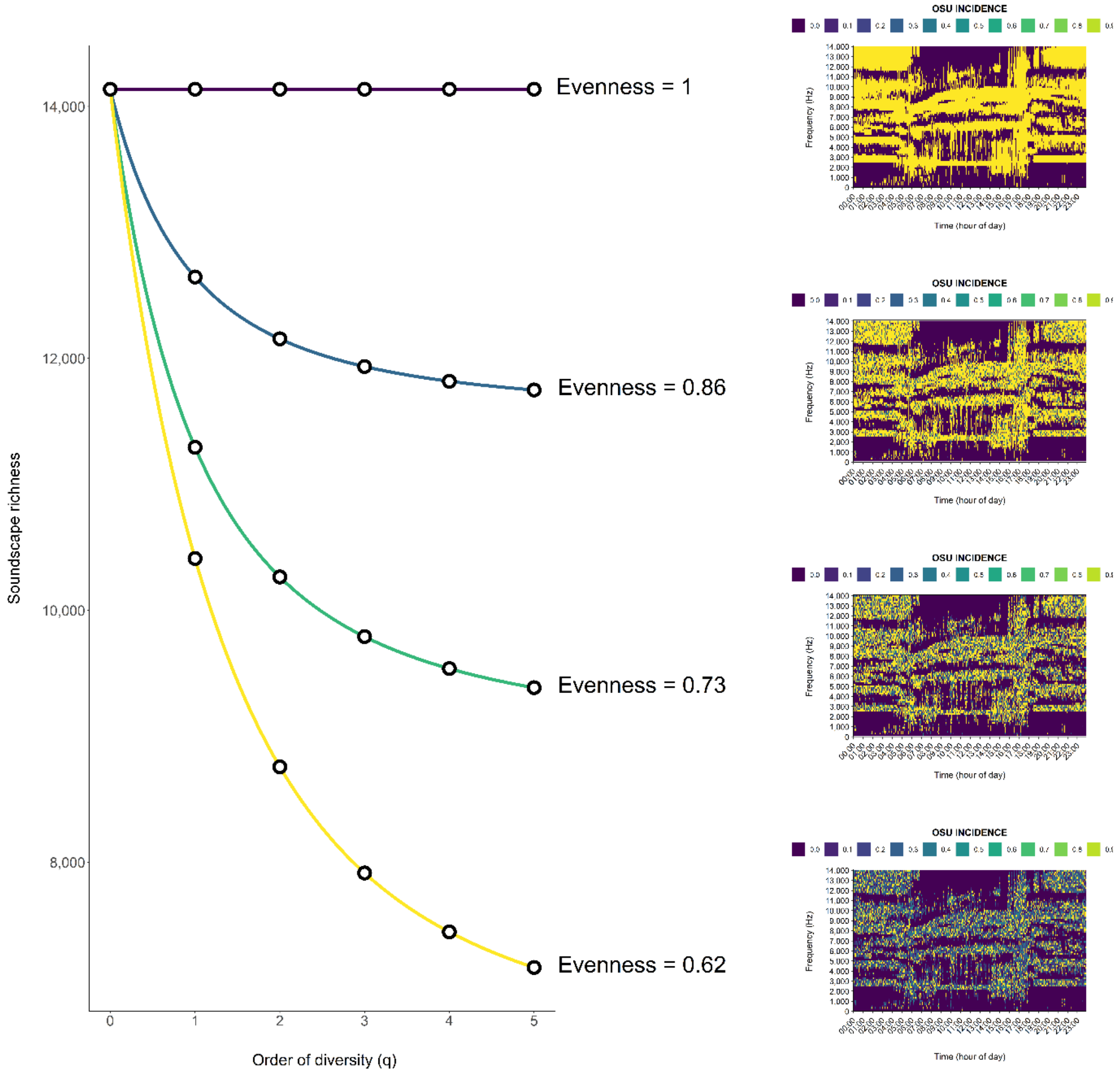


***Figure S11:*** *A soundscape diversity profile displaying the change in the soundscape diversity in function of the order of diversity (q) for four soundscapes with differing evenness ($^0D/ {}^2D$) values. The lower the evenness of the soundscape, the more rapidly the soundscape diversity drops in function of the order of diversity.*

### 5.6. Soundscape diversity can be decomposed into its alpha, beta, and gamma components

The soundscape diversity can be assessed with reference to the wider ecological system by breaking it down into its respective alpha, beta and gamma components (Whittaker 1960). Within the framework of Hill numbers, these components take a simple multiplicative relationship in which gamma equals alpha times beta.

To illustrate the meaning of these components, in the following example (Fig. S12) we computed the alpha, beta and gamma components for three hypothetical scenarios: (i) a system comprised of two identical and equally diverse soundscapes (Fig. S12A); (ii) a system comprised of two unique but equally diverse soundscapes (Fig. S12B); (iii) a system comprised of two partially overlapping soundscapes (Fig. S12C). To create these hypothetical scenarios, we used the same artificial soundscapes generated in section S.5.4. For the first scenario, we replicated the same soundscape twice (soundscape 1 and soundscape 1), and computed the alpha, beta, and gamma components for the system. For the second scenario, we took the two equally diverse soundscapes with no common OSUs (soundscape 1 and soundscape 2) and computed the three components. Finally, for the third scenario, we computed alpha, beta and gamma for a system comprised of soundscape 1 and the pooled soundscape, which partially overlaps the former.

When two soundscapes were identical, alpha equalled gamma and thus beta equalled 1. When two soundscapes were equally diverse but unique, gamma was double alpha, and thus beta equalled 2. When two soundscapes overlapped partially, beta ranged between 1 and N (the total number of soundscapes). In this case, beta could be seen as the effective number of equally large and completely distinct soundscapes in the overall system – or a measure of how many times more diverse the whole system (gamma) was in the effective number of OSUs compared to its soundscapes on average.

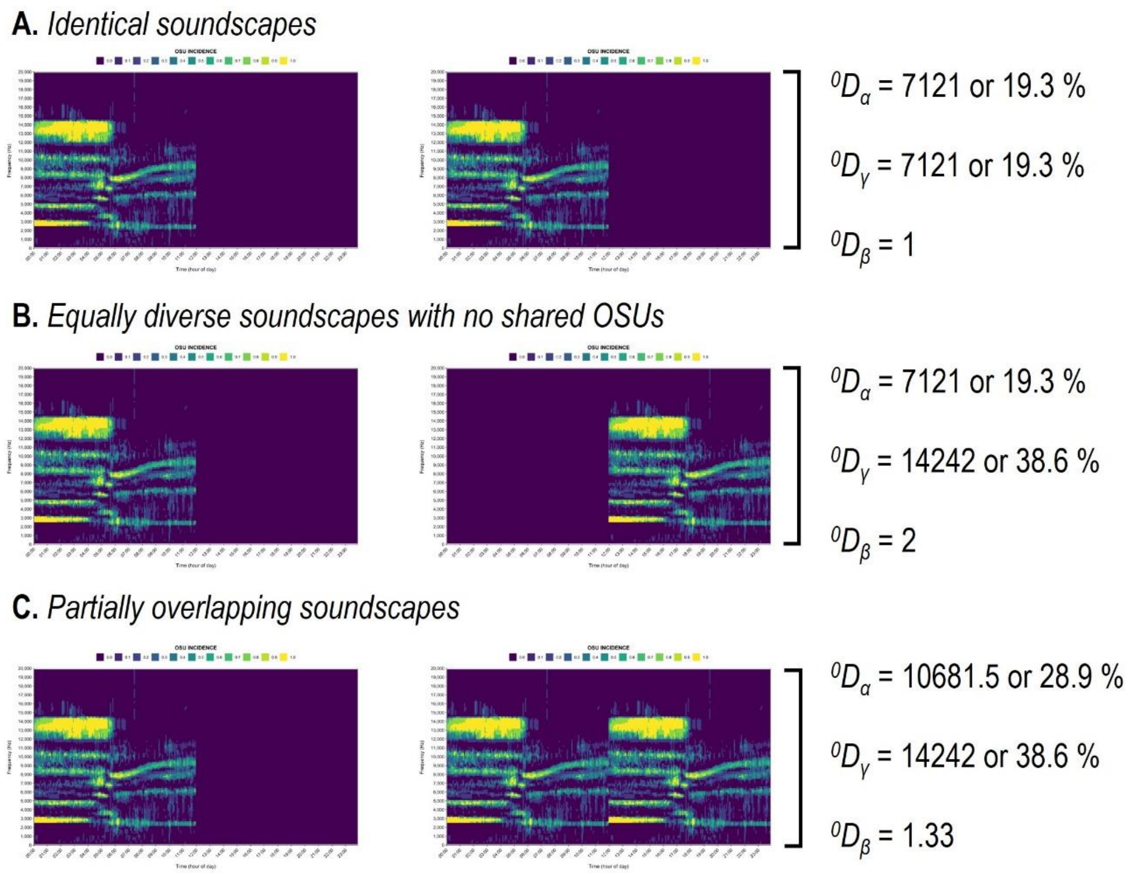


***Figure S12:*** *A simple example illustrating soundscape diversity partitioning for three scenarios:* ***A.*** *Two equally diverse and completely identical soundscapes with the same relative abundance distribution. In this scenario, gamma equals alpha, and beta – or the number of equally large and completely distinct soundscapes – is one.* ***B.*** *Two equally diverse but unique soundscapes. In this scenario, gamma is twice alpha, and the number of distinct soundscapes (beta) is 2 – or the number of subsystems.* ***C.*** *Two partially overlapping soundscapes, one of which is twice as diverse as the other. In this scenario, beta ranges between 1 and N (the number of soundscapes).*

### S.6. Additional case study information

#### 6.1. Site selection

The case study described in the main text used the set of recordings detailed in Bueno et al. (2020), collected at 151 plots at 78 sites (74 forest islands and four continuous forest sites) at the Balbina Hydroelectric Reservoir (1°40'S, 59°40'W) in Central Amazonia. For the purposes of the case study, only the plots for which data on the richness of soniferous species was available were retained (see section 6.2. for further details). Moreover, all plots located in riparian habitats near streams were removed from the dataset. Additionally, the sound recordings for all plots were visually inspected using long-duration false-colour spectrograms (LDFCSs) (see Towsey et al. 2016; Towsey et al. 2018), and plots with signs of microphone failure or persistent noise were removed. Ultimately, 65 plots located at 35 sites (32 islands and three continuous forest sites) were retained.

***Table S3:*** *Overview of the sites and their respective sub-plots included in the case study*

| Island name | Plot name | Island name | Plot name |
|---|---|---|---|
| Abusado | Abusado | Jabuti | Jabuti_A |
| Adeus | Adeus_A | | Jabuti_B |
| | Adeus_B | | Jabuti_C |
| Aline | Aline | Jiquitaia | Jiquitaia |
| Andre | Andre | Joaninha | Joaninha |
| Arrepiado | Arrepiado | Martelo | Martelo_B |
| Bacaba | Bacaba_B | | Martelo_C |
| Beco_do_Catitu | Beco_do_Catitu_A | Mascote | Mascote_A1 |
| | Beco_do_Catitu_B | | Mascote_A2 |
| | Beco_do_Catitu_D | | Mascote_B1 |
| | Beco_do_Catitu_E | | Mascote_B2 |
| Cafundo | Cafundo | Moita | Moita_A |
| CF_Grid | CF_Grid_CampTrail_A | | Moita_B |
| | CF_Grid_NS3_1200 | Palhal | Palhal |
| CF_Loreno | CF_Loreno_A | Panema | Panema |
| | CF_Loreno_B | Pe_Torto | Pe_Torto |
| CF_WABA | CF_WABA_B | Piquia | Piquia |
| | CF_WABA_C | Pontal | Pontal_B |
| Cipoal | Cipoal_A | | Pontal_C |
| | Cipoal_B | Porto_Seguro | Porto_Seguro_B |
| | Cipoal_C | | Porto_Seguro_C |
| Coata | Coata | | Porto_Seguro_D |
| Formiga | Formiga | Relogio | Relogio_B |
| Furo_de_Santa_Luzia | Furo_de_Santa_Luzia_B | Sapupara | Sapupara_A |
| | Furo_de_Santa_Luzia_C | | Sapupara_B |
| Fuzaca | Fuzaca_B | Torem | Torem |
| | Fuzaca_C | Tristeza | Tristeza_A |
| | Fuzaca_D | | Tristeza_B |
| Garrafa | Garrafa | | Tristeza_C |
| Gaviao_real | Gaviao_real_A | Tucumari | Tucumari_A |
| | Gaviao_real_B | | Tucumari_B |
| | Gaviao_real_C | | Tucumari_C |
| | Gaviao_real_D | | |

### 6.2. Compound richness of soniferous species

To assess the relationship between the soundscape diversity metrics and the taxonomic richness of sound-producing organisms in the study area, we calculated the compound richness of three major vocalising groups in tropical rain forests: (i) anurans; (ii) birds; and (iii) primates. Below, we outlined how the taxonomic richness for each of these groups was determined.

#### 6.2.1. Anuran data

The anuran species richness was determined using passive acoustic monitoring data obtained at BHR. Per plot, a subset of 62 1-min recordings was selected for aural identification of anuran species, taking the first 1-min of recording every 10-min over 5 hours between 17:00h-22:01h during sampling days 2 and 4. In each sound recording, anuran species were identified using both aural identification and visual inspection of spectrograms by a trained expert (GSM) using the RFCx Arbimon II Visualizer Tool (https://arbimon.rfcx.org/). All species identifications were cross-validated by a second reviewer (ILK) to ensure accuracy, and species records were discarded if they could not be readily identified or if spectrograms were inadequate.

#### 6.2.2. Bird data

The detection history of species in the audio recordings was acquired through three steps. First, an expert (MC-C) manually searched for species in recordings from two non-consecutive days in each plot and created a preliminary list of species and a call template for each species detected. Second, in the RFCx-ARBIMON platform, we used the template matching procedure to classify the audio recordings. In this step, we created two audio playlists: a) all diurnal (05:00h - 18:00h) recordings and b) all nocturnal (18:00h - 05:00h) recordings. All bird templates, except for *Glaucidium hardyi* were assigned to the nocturnal playlist. The classification of recordings based on the pattern matching procedures searches through the audio data (~ X 1-min recordings) for acoustic signals and detects regions that have a high correlation with a template that has been selected by the user. All regions of interest (ROIs) with values above the selected correlation threshold (0.1) were presented as potential detections for posterior validations (LeBien et al. 2020). We then used a filter to display only the best matches per plot per day. The selection of a low threshold resulted in a high number of false positives, though the number of false negatives was negligible. In the third step, one of us (MC-C) manually reviewed the template matching results using a filter that displays only the best matches per plot and per day. In this step, we annotated the results as either positive or negative, indicating the corresponding species presence or absence. This ensured that the final data set used in the analyses only included expert verified detections and the exclusion of all false positive detections.

#### 6.2.3. Primate data

The primate data was compiled from Benchimol and Peres (2015). In this study, vertebrate surveys including primates were conducted on 37 islands and 3 continuous forest sites. Primate surveys were conducted based on diurnal line-transect censuses, although some species were also detected using a systematic camera-trapping programme (Benchimol and Peres 2021). For each of the sites, one to five line-transects of variable length were cut based on the island size and shape. For the continuous forest sites, three parallel 4-km transects were established with a 1-km separation. At each of the sites, line-transect surveys were conducted by expert observers, walking the transects at a constant speed (~ 1 km/h) in the morning (06:15h - 10:30h) and afternoon (14:00h - 17:30h) following a standard protocol (Peres and Cunha 2011). For each sampling year of the study (2011 and 2012), four line-transect surveys were conducted per site, each of which was separated by 30 sampling days to minimize the impact of time of day and seasonality.

In addition to the line-transect surveys, sign surveys for vertebrate activity were conducted on the return walks, and species identifications were recorded. Moreover, to supplement these surveys, Reconyx HC 500 Hyperfire camera traps were deployed at each site. For each sampling site, two to ten camera traps were deployed at 30-40 cm height, separating individual camera traps at an approximate distance of 500 m depending on island size and shape. For the continuous forest sites, 15 camera traps were deployed, installing five camera traps per transect. All camera trap stations were active for 30 days in each sampling year, taking a sequence of five photos for every detection event, and using 15-second intervals between consecutive detections. Camera trapping efforts were always temporally separated from line-transect surveys to prevent disturbance of the local fauna. All data from line-transect surveys, sign surveys and camera trapping were compiled into presence-absence data per island for the species known to present in the study area.

#### 6.2.4. Compound species richness

A subset of the species richness data was taken, including only the sites and plots outlined in section 6.1 (Table S3). For sampling sites containing multiple plots, the site-wide species richness was calculated for each taxonomic group. Finally, for each site, the richness values for the anurans, birds and primates were summed to obtain the compound richness of soniferous species. This metric of taxonomic richness was compared against the soundscape diversity metrics to assess their behaviour along this ecological gradient (Table S4).

***Table S4:*** *Soundscape diversity and taxonomic richness data for 35 sites at the Balbina Hydroelectric Reservoir*

| **Site** | **Soundscape diversity data** | | **Taxonomic richness data** | | | |
|---|---|---|---|---|---|---|
| | soundscape richness (%) | soundscape evenness | anuran richness | bird richness | primate richness | compound richness |
| Abusado | 30.44 | 0.66 | 9 | 4 | 3 | 16 |
| Adeus | 53.52 | 0.68 | 7 | 15 | 2 | 24 |
| Aline | 36.13 | 0.61 | 4 | 6 | 0 | 10 |
| Andre | 23.18 | 0.61 | 9 | 3 | 1 | 13 |
| Arrepiado | 27.27 | 0.63 | 5 | 10 | 1 | 16 |
| Bacaba | 40.83 | 0.66 | 4 | 10 | 4 | 18 |
| Beco_do_Catitu | 59.80 | 0.70 | 17 | 20 | 6 | 43 |
| Cafundo | 37.26 | 0.77 | 7 | 10 | 0 | 17 |
| CF_Grid | 56.23 | 0.73 | 12 | 21 | 7 | 40 |
| CF_Loreno | 61.93 | 0.72 | 10 | 21 | 7 | 38 |
| CF_Waba | 60.35 | 0.67 | 12 | 16 | 7 | 35 |
| Cipoal | 57.80 | 0.68 | 13 | 25 | 7 | 45 |
| Coata | 42.37 | 0.63 | 10 | 15 | 2 | 27 |
| Formiga | 33.15 | 0.66 | 7 | 3 | 0 | 10 |
| Furo_de_Santa_Luzia | 55.71 | 0.68 | 7 | 17 | 5 | 29 |
| Fuzaca | 53.24 | 0.72 | 16 | 18 | 7 | 41 |
| Garrafa | 17.68 | 0.63 | 6 | 6 | 1 | 13 |
| Gaviao_real | 44.43 | 0.70 | 13 | 24 | 5 | 42 |
| Jabuti | 63.40 | 0.72 | 11 | 21 | 7 | 39 |
| Jiquitaia | 36.09 | 0.60 | 1 | 4 | 3 | 8 |
| Joaninha | 33.96 | 0.62 | 4 | 3 | 0 | 7 |
| Martelo | 56.79 | 0.69 | 10 | 21 | 7 | 38 |
| Mascote | 66.50 | 0.72 | 13 | 28 | 7 | 48 |
| Moita | 47.05 | 0.66 | 7 | 17 | 4 | 28 |
| Palhal | 33.29 | 0.65 | 3 | 6 | 2 | 11 |
| Panema | 35.80 | 0.77 | 3 | 2 | 0 | 5 |
| Pe_Torto | 34.66 | 0.65 | 3 | 7 | 1 | 11 |
| Piquia | 39.69 | 0.65 | 7 | 9 | 1 | 17 |
| Pontal | 48.15 | 0.66 | 10 | 21 | 5 | 36 |
| Porto_Seguro | 58.69 | 0.71 | 21 | 27 | 7 | 55 |
| Relogio | 52.86 | 0.65 | 8 | 8 | 6 | 22 |
| Sapupara | 57.06 | 0.70 | 10 | 11 | 5 | 26 |
| Torem | 27.43 | 0.79 | 3 | 2 | 1 | 6 |
| Tristeza | 66.02 | 0.74 | 13 | 23 | 7 | 43 |
| Tucumari | 50.44 | 0.67 | 9 | 26 | 4 | 39 |

### Publication bibliography


Aide, T.; Hernández-Serna, Andres; Campos-Cerqueira, Marconi; Acevedo-Charry, Orlando; Deichmann, Jessica (2017): Species Richness (of Insects) Drives the Use of Acoustic Space in the Tropics. In *Remote Sensing* 9 (11), p. 1096. DOI: 10.3390/rs9111096.

Alberdi, Antton; Gilbert, M. Thomas P. (2019): A guide to the application of Hill numbers to DNA-based diversity analyses. In *Mol Ecol Resour* 19 (4), pp. 804–817. DOI: 10.1111/1755-0998.13014.

Baty, Florent; Ritz, Christian; Charles, Sandrine; Brutsche, Martin; Flandrois, Jean-Pierre; Delignette-Muller, Marie-Laure; others (2015): A toolbox for nonlinear regression in R: the package nlstools. In *Journal of Statistical Software* 66 (5), pp. 1–21.

Benchimol, Maíra; Peres, Carlos A. (2015): Widespread Forest Vertebrate Extinctions Induced by a Mega Hydroelectric Dam in Lowland Amazonia. In *PLOS ONE* 10 (7), e0129818. DOI: 10.1371/journal.pone.0129818.

Benchimol, Maíra; Peres, Carlos A. (2021): Determinants of population persistence and abundance of terrestrial and arboreal vertebrates stranded in tropical forest land-bridge islands. In *Conservation Biology* 35 (3), pp. 870–883. DOI: 10.1111/cobi.13619.

Bradfer-Lawrence, Tom; Gardner, Nick; Bunnefeld, Lynsey; Bunnefeld, Nils; Willis, Stephen G.; Dent, Daisy H. (2019): Guidelines for the use of acoustic indices in environmental research. In *Methods Ecol Evol* 10 (10), pp. 1796–1807. DOI: 10.1111/2041-210X.13254.

Bueno, Anderson Saldanha; Masseli, Gabriel S.; Kaefer, Igor L.; Peres, Carlos A. (2020): Sampling design may obscure species–area relationships in landscape-scale field studies. In *Ecography* 43 (1), pp. 107–118. DOI: 10.1111/ecog.04568.

Burivalova, Zuzana; Towsey, Michael; Boucher, Tim; Truskinger, Anthony; Apelis, Cosmas; Roe, Paul; Game, Edward T. (2018): Using soundscapes to detect variable degrees of human influence on tropical forests in Papua New Guinea. In *Conservation biology : the journal of the Society for Conservation Biology* 32 (1), pp. 205–215. DOI: 10.1111/cobi.12968.

Campos-Cerqueira, Marconi; Mena, Jose Luis; Tejeda-Gómez, Vania; Aguilar-Amuchastegui, Naikoa; Gutierrez, Nelson; Aide, T. Mitchell (2020): How does FSC forest certification affect the acoustically active fauna in Madre de Dios, Peru? In *Remote Sens Ecol Conserv* 6 (3), pp. 274–285. DOI: 10.1002/rse2.120.

Chao, Anne; Chiu, Chun-Huo; Hsieh, T. C. (2012): Proposing a resolution to debates on diversity partitioning. In *Ecology* 93 (9), pp. 2037–2051. DOI: 10.1890/11-1817.1.

Chao, Anne; Chiu, Chun-Huo; Jost, Lou (2014): Unifying Species Diversity, Phylogenetic Diversity, Functional Diversity, and Related Similarity and Differentiation Measures Through Hill Numbers. In *Annu. Rev. Ecol. Evol. Syst.* 45 (1), pp. 297–324. DOI: 10.1146/annurev-ecolsys-120213-091540.

Chao, Anne; Jost, Lou (2012): Coverage-based rarefaction and extrapolation: standardizing samples by completeness rather than size. In *Ecology* 93 (12), pp. 2533–2547. DOI: 10.1890/11-1952.1.

Chao, Anne; Jost, Lou (2015): Estimating diversity and entropy profiles via discovery rates of new species. In *Methods Ecol Evol* 6 (8), pp. 873–882. DOI: 10.1111/2041-210X.12349.

Chiu, Chun-Huo; Jost, Lou; Chao, Anne (2014): Phylogenetic beta diversity, similarity, and differentiation measures based on Hill numbers. In *Ecological Monographs* 84 (1), pp. 21–44. DOI: 10.1890/12-0960.1.

Eldridge, Alice; Guyot, Patrice; Moscoso, Paola; Johnston, Alison; Eyre-Walker, Ying; Peck, Mika (2018): Sounding out ecoacoustic metrics: Avian species richness is predicted by acoustic indices in temperate but not tropical habitats. In *Ecological Indicators* 95, pp. 939–952. DOI: 10.1016/j.ecolind.2018.06.012.

Gasc, A.; Sueur, J.; Jiguet, F.; Devictor, V.; Grandcolas, P.; Burrow, C. et al. (2013): Assessing biodiversity with sound: Do acoustic diversity indices reflect phylogenetic and functional diversities of bird communities? In *Ecological Indicators* 25, pp. 279–287. DOI: 10.1016/j.ecolind.2012.10.009.

Hallett, Lauren M.; Jones, Sydney K.; MacDonald, A. Andrew M.; Jones, Matthew B.; Flynn, Dan F. B.; Ripplinger, Julie et al. (2016): codyn : An r package of community dynamics metrics. In *Methods Ecol Evol* 7 (10), pp. 1146–1151. DOI: 10.1111/2041-210X.12569.

Harrison, Susan; Ross, Sally J.; Lawton, John H. (1992): Beta Diversity on Geographic Gradients in Britain. In *The Journal of Animal Ecology* 61 (1), p. 151. DOI: 10.2307/5518.

Hill, M. O. (1973): Diversity and Evenness: A Unifying Notation and Its Consequences. In *Ecology* 54 (2), pp. 427–432. DOI: 10.2307/1934352.

Hsieh, T. C.; Ma, K. H.; Chao, Anne (2016): iNEXT: an R package for rarefaction and extrapolation of species diversity (H ill numbers). In *Methods Ecol Evol* 7 (12), pp. 1451–1456. DOI: 10.1111/2041-210X.12613.

Jost, Lou (2006): Entropy and diversity. In *Oikos* 113 (2), pp. 363–375. DOI: 10.1111/j.2006.0030-1299.14714.x.

Jost, Lou (2007): Partitioning diversity into independent alpha and beta components. In *Ecology* 88 (10), pp. 2427–2439. DOI: 10.1890/06-1736.1.

Landini, G.; Randell, D. A.; Fouad, S.; Galton, A. (2017): Automatic thresholding from the gradients of region boundaries. In *Journal of microscopy* 265 (2), pp. 185–195. DOI: 10.1111/jmi.12474.

LeBien, Jack; Zhong, Ming; Campos-Cerqueira, Marconi; Velev, Julian P.; Dodhia, Rahul; Ferres, Juan Lavista; Aide, T. Mitchell (2020): A pipeline for identification of bird and frog species in tropical soundscape recordings using a convolutional neural network. In *Ecological Informatics* 59, p. 101113. DOI: 10.1016/j.ecoinf.2020.101113.

Leinster, Tom; Cobbold, Christina A. (2012): Measuring diversity: The importance of species similarity. In *Ecology* 93 (3), pp. 477–489. DOI: 10.1890/10-2402.1.

Machado, Ricardo B.; Aguiar, Ludmilla; Jones, Gareth (2017): Do acoustic indices reflect the characteristics of bird communities in the savannas of Central Brazil? In *Landscape and Urban Planning* 162, pp. 36–43. DOI: 10.1016/j.landurbplan.2017.01.014.

Mouchet, Maud A.; Villéger, Sébastien; Mason, Norman W. H.; Mouillot, David (2010): Functional diversity measures: an overview of their redundancy and their ability to discriminate community assembly rules. In *Functional Ecology* 24 (4), pp. 867–876. DOI: 10.1111/j.1365-2435.2010.01695.x.

Peres, C. A.; Cunha, A. A. (2011): Manual para censo e monitoramento de vertebrados de médio e grande porte por transecção linear em florestas tropicais. In *Wildlife Conservation Society, Brasılia, Brasil*.

Phillips, Yvonne F.; Towsey, Michael; Roe, Paul (2018): Revealing the ecological content of long-duration audio-recordings of the environment through clustering and visualisation. In *PloS one* 13 (3), e0193345. DOI: 10.1371/journal.pone.0193345.

R Core Team (2020): R: A Language and Environment for Statistical Computing. Vienna, Austria. Available online at https://www.r-project.org/.

Ricotta, Carlo (2005): A note on functional diversity measures. In *Basic and Applied Ecology* 6 (5), pp. 479–486. DOI: 10.1016/j.baae.2005.02.008.

Rodriguez, Alexandra; Gasc, Amandine; Pavoine, Sandrine; Grandcolas, Philippe; Gaucher, Philippe; Sueur, Jérôme (2014): Temporal and spatial variability of animal sound within a neotropical forest. In *Ecological Informatics* 21, pp. 133–143. DOI: 10.1016/j.ecoinf.2013.12.006.

Towsey, Michael; Truskinger, Anthony; Roe, Paul (2016): QUT Ecoacoustics | Long-duration Audio-recordings of the Environment. QUT Ecoacoustics Group. Available online at https://research.ecosounds.org/research/eadm-towsey/long-duration-audio-recordings-of-the-environment, updated on 1/13/2021, checked on 1/13/2021.

Towsey, Michael; Znidersic, Elizabeth; Broken-Brow, Julie; Indraswari, Karlina; Watson, David M.; Phillips, Yvonne et al. (2018): Long-duration, false-colour spectrograms for detecting species in large audio data-sets. In *JEA* 2 (1), p. 1. DOI: 10.22261/JEA.IUSWUI.

Tuomisto, Hanna (2010): A diversity of beta diversities: Straightening up a concept gone awry. Part 1. Defining beta diversity as a function of alpha and gamma diversity. In *Ecography* 33 (1), pp. 2–22. DOI: 10.1111/j.1600-0587.2009.05880.x.

Villéger, Sébastien; Mason, Norman W. H.; Mouillot, David (2008): New multidimensional functional diversity indices for a multifaceted framework in functional ecology. In *Ecology* 89 (8), pp. 2290–2301. DOI: 10.1890/07-1206.1.

Whittaker, R. H. (1960): Vegetation of the Siskiyou Mountains, Oregon and California. In *Ecological Monographs* 30 (3), pp. 279–338. DOI: 10.2307/1943563.
